# Relaxation time asymmetry in stator dynamics of the bacterial flagellar motor

**DOI:** 10.1101/2021.07.05.451114

**Authors:** Ruben Perez-Carrasco, María-José Franco-Oñate, Jean-Charles Walter, Jérôme Dorignac, Fred Geniet, John Palmeri, Andrea Parmeggiani, Nils-Ole Walliser, Ashley L Nord

**Affiliations:** Department of Life Sciences, Imperial College London, London, SW7 2BU UK; Laboratoire Charles Coulomb (L2C), Univ. Montpellier, CNRS, Montpellier, France; Centre de Biologie Structurale U. Montpellier, CNRS, INSERM, Montpellier, 34090 France

## Abstract

The bacterial flagellar motor (BFM) is the membrane-embedded rotary molecular motor which turns the flagellum that provides thrust to many bacterial species. This large multimeric complex, composed of a few dozen constituent proteins, has emerged as a hallmark of dynamic subunit exchange. The stator units are inner-membrane ion channels which dynamically bind and unbind to the peptidoglycan at the rotor periphery, consuming the ion motive force (IMF) and applying torque to the rotor when bound. The dynamic exchange is known to be a function of the viscous load on the flagellum, allowing the bacterium to dynamically adapt to its local viscous environment, but the molecular mechanisms of exchange and mechanosensitivity remain to be revealed. Here, by actively perturbing the steady-state stator stoichiometry of individual motors, we reveal a stoichiometry-dependent asymmetry in stator remodeling kinetics. We interrogate the potential effect of next-neighbor interactions and local stator unit depletion and find that neither can explain the observed asymmetry. We then simulate and fit two mechanistically diverse models which recapitulate the asymmetry, finding stator assembly dynamics to be particularly well described by a two-state catch-bond mechanism.

---

The bacterial flagellar motor (BFM) is the rotary molecular motor which, in many bacterial species, provides the thrust necessary for motility, chemotaxis, biofilm formation and infection. The BFM is large (∼ 11 MDa), dynamically self-assembled, and membrane-spanning, comprised of more than a dozen different proteins. The rotor, which extends into the cytoplasm, is rotated by up to around a dozen peptidogylcan (PG)-bound stator units, each one powered by the electrochemical gradient across the cell membrane. This rotation is coupled to the extracellular flagellar filament, which can spin at speeds of up to hundreds of Hertz.

Historically, it was imagined that, once assembled, the composition of the BFM was static. However, thanks to a multitude of studies over the last decade showing dynamic exchange of multiple motor components, the BFM has become a hallmark of dynamic subunit exchange in multimeric protein complexes [1]. This phenomenon has been most well characterized for the stator units, which exchange between an active, motor-bound, torque-producing population and a pool of inactive membrane-diffusing units [2]. While the purpose and mechanism of this exchange have not yet been fully elucidated, the stator units are mechanosensitive, with larger viscous loads on the flagellum leading to higher stator occupancy [3, 4]. Stator unit exchange occurs on timescales of seconds to minutes, allowing the motors of individual cells to quickly and dynamically adapt to changes in their environment.

The dynamics of stator remodeling was first quantitatively modeled with a simple reversible Hill-Langmuir adsorption model [2, 5]. When this model was applied to motors driving various viscous loads, it was observed that the stator unbinding rate was load-dependent, and that the stator units behaved in a manner consistent with a catch-bond mechanism. In spite of these results, the particular mechanism in control of the stoichiometry dynamics remains elusive. Recently, Wadwha et al. proposed a more complex model which conjectures that, in addition to a load-dependent unbinding rate, both the binding and unbinding rates are also dependent upon the speed of the motor [6]. In contrast to the Hill-Langmuir model, this model presents an intriguing and testable prediction: the characteristic relaxation timescale towards steady state depends on the initial stator stoichiometry. Here, by employing a variety of experimental methods, we explore different starting stoichiometries of individual motors and measure their relaxation dynamics. In order to shed light on the mechanistic nature of the resulting timescales, we compare the experimental results with the model of Wadwha et al. and with other models compatible with a relaxation time depending on the initial stoichiometry. In particular, we propose an alternative model based on a two-state catch bond, and we discuss the implications of these results with respect to stator remodeling of the BFM and more generally for dynamic protein complexes.

## Results

### Measurements of BFM relaxation to steady state

Previously, we used magnetic tweezers to reversibly increase the viscous load on individual motors, transiently perturbing the stator stoichiometry from steady state, then quantified stator dynamics during the relaxation back to steady state [5]. In this work, we started by repeating these measurements. Cells of a non-switching strain of *E. coli* lacking flagellar filaments were immobilized to a coverslip, and streptavidin-coated superparamagnetic particles were attached to an endogenously biotinylated hook. The position of the particles was tracked and used as a proxy for the angular position of the motor, yielding motor speed and torque. Following a steady-state measurement, two permanent magnets were brought near the sample. Here, the torque exerted by the magnets on the particle counters and balances the torque exerted by the BFM upon the particle, thereby stalling motor rotation. This resulted in the recruitment of additional stators, priming the system to a state of higher stoichiometry than that of steady state. Following 10 minutes of motor stall, the magnets were rapidly removed, and the speed and torque of the motor were measured as the system relaxed back to steady state.

Following relaxation, and using the exact same set of motors, we perturbed the motors away from steady-state stoichiometry, but in the opposite direction, towards lower stator stoichiometries. We introduced an ionophore to collapse the proton motive force (PMF) and waited 8 min for the stators to fully dissociate from the motor. We then flushed out the ionophore and measured motor speed and torque as the stators re-incorporated to reach steady state, a process we refer to as ‘resurrection’. All measurements were performed for three viscous loads (*γ*_1300_, *γ*_500_, *γ*_300_) by using beads of 1300, 500, and 300 nm in diameter. Details are given in Materials and Methods and as described previously [5].

From these different experiments we obtained, for each individual motor, traces in time for three different initial stoichiometry conditions: steady state, release from stall, and resurrection. Using a step detection algorithm and knowledge of the torque per stator from individual traces, we calculated stator stoichiometry trajectories in time [5]. Fig. 1A shows an example of a single motor measurement, Fig. 1B shows a schematic of the evolution of stoichiometry, and Fig. 1C-D shows motor torque and stator stoichiometry as a function of time for all of the measurements.

**Figure 1:**
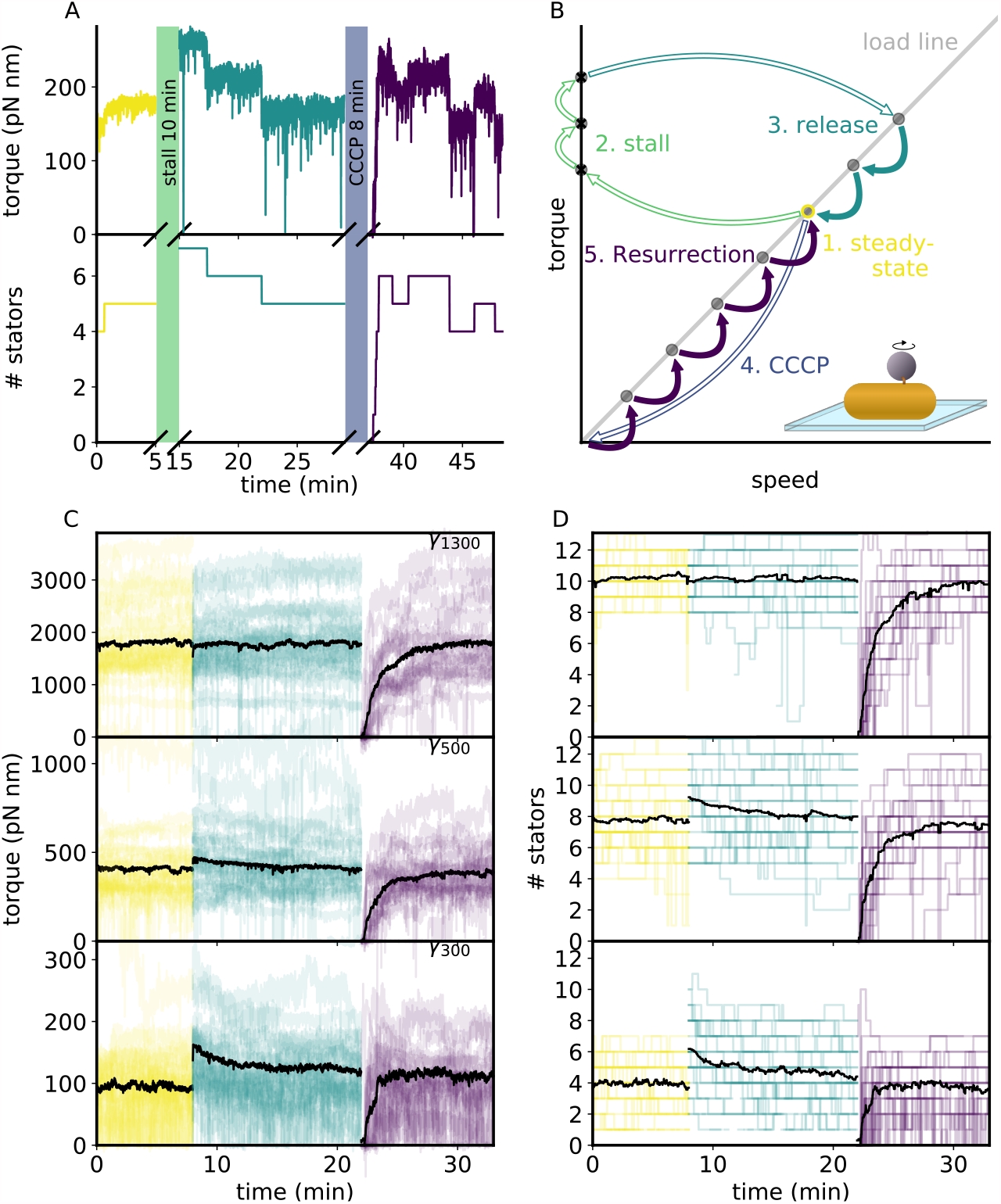
A) Single measurement of a 300 nm bead, showing motor torque and stator number as a function of time. The experimental protocol for single motor measurements was the following: unperturbed steady-state measurement (yellow), motor stall via magnetic tweezers (10 min), release from stall and relaxation back to steady state (green), introduction of ionophore (8 min), removal of ionophpre and motor resurrection to steady state (purple). B) A schematic of the torque-speed plane for the example in (A), where colors correspond to the experimental steps. Solid arrows represent observed transitions, open arrows non-observed transitions. Bottom inset shows the experimental assay. C) Individual traces of motor torque versus time and D) stator number versus time for the steady state (yellow), release from stall (green), and resurrection (purple) measurements. Black lines show the average behavior of all motors. Subplots from top to bottom shows 1300, 500, and 300 nm beads (27, 33, and 31 measurements, respectively).

### An asymmetry in the characteristic relaxation time

We previously proposed a reversible Hill-Langmuir adsorption model to describe stator assembly kinetics [5]. In this model, the rotor is surrounded by *N*_max_ fixed, independent, and non-interacting binding sites. Stator units freely diffusing in the inner membrane can bind to an empty site on the rotor with rate constant *k*_on_, bound stator units can dissociate with rate constant *k*_off_, and the average number of bound stator units, ⟨*N*⟩, evolves according to

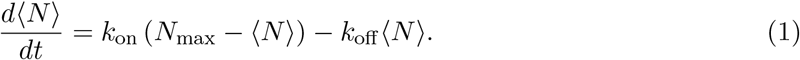

We assume the concentration of freely diffusing stators is large enough to be considered constant. The analytical solution,

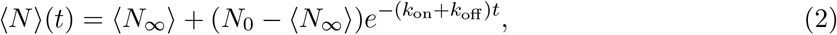

relaxes from stator occupancy *N*_0_ at *t* = 0 to the average steady-state value, ⟨*N*_*∞*_⟩ = *N*_max_*/*(1 + *k*_off_ */k*_on_), with a characteristic relaxation time *t*_*c*_ ≡ 1*/*(*k*_on_ + *k*_off_).

Eq. 2 was fit to the average over all motors for each experimental relaxation (shown in Fig. 2A), with the exception of release from stall for *γ*_1300_, wherein the number of stators is not significantly affected by motor stall. The resulting rate constants and characteristic relaxation times are shown in Fig. 2B. In accordance with earlier work, we observe that *t*_*c*_ is dependent upon load, and thus upon the torque delivered by a single stator unit: the smaller the single stator unit torque, the faster the relaxation. Unexpectedly, we observed a new feature which becomes apparent only when the system is prepared into different starting stoichiometries: the timescale to approach steady state is asymmetric; resurrection traces are faster than release from stall traces. This asymmetry (dependence of *t*_*c*_ on the starting stoichiometry) is not compatible with the previously proposed Hill-Langmuir adsorption model, and opens the door to understanding the mechanisms that control the feedback between load and stoichiometry dynamics. Consequently, we proceeded by interrogating various potential mechanisms in search of the source of this observed asymmetry.

**Figure 2:**
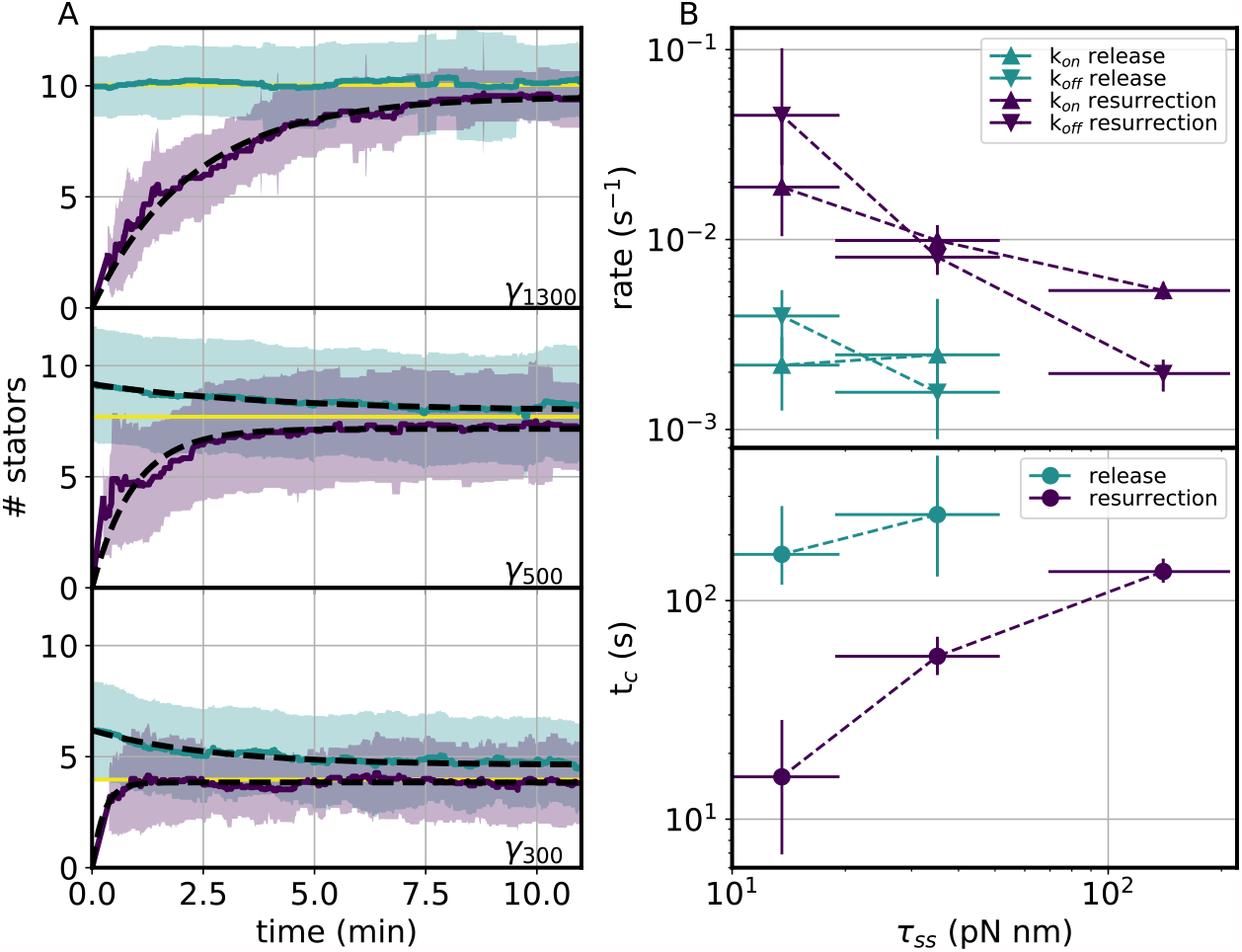
A) Evolution of stator stoichiometry for release from stall (green) and resurrection (purple). Viscous loads, from the top to bottom are *γ*_1300_, *γ*_500_, and *γ*_300_. Green and purple lines show the average of multiple traces, shading shows standard deviation, and the yellow line shows the average of steady-state measurements. The black dashed line shows the best fit of Eq. 2 to each time series (with the exception of release from stall for *γ*_1300_). B) The rates (top) and characteristic times (bottom) as a function of single stator torque at steady state, *τ*_*ss*_. Error bars on *τ*_*ss*_ are the standard deviation of all measurements. Error bars on rates and *t*_*c*_ represent the 90% high density interval from Approximate Bayesian Computation inference.

### Neither depletion nor cooperativity explain the observed asymmetry

We investigated whether either next-neighbor unit-unit interactions or local depletion of unbound stator units surrounding the motor could explain the observed asymmetry in relaxation time. We first employed a grand canonical Hamiltonian description of a one-dimensional Short Range Lattice Gas (SRLG) with periodic boundary conditions to rigorously explore the effects of stator cooperativity (see SI for more details). Our Glauber Monte Carlo simulations based on this model show that the characteristic relaxation time increases quickly with the (dimensionless) unit-unit interaction energy *βJ* ≥ 0 (where *β* = (*k*_B_*T*)^*−*1^, with the Boltzmann constant *k*_*B*_ and T the temperature), and the correlation length, the characteristic distance over which the occupancy of one binding site influences that of its neighbors, grows as well [7–9] (Fig. 1). Furthermore, the characteristic relaxation time for positive values of *βJ* depends on the average occupancy at steady state ⟨*N*_*∞*_⟩. Note that when the (dimensionless) interaction energy *βJ* equals zero, the SRLG is equivalent to the Hill-Langmuir adsorption model, and the following relation between the effective (dimensionless) chemical potential of the reservoir, 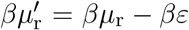 and the Hill-Langmuir rate constants is satisfied: *βµ*_r_ = log (*k*_on_*/k*_off_). Here, *µ*_r_ is the chemical potential of the reservoir, *ε* is the binding energy.

Simulation results do indeed exhibit different relaxation times to steady state between release from stall and resurrection, 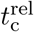 and 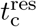, respectively. The cooperativity-induced time asymmetry cannot, however, reasonably explain the experimentally measured ones, even though the former increases with the nearest-neighbor (dimensionless) interaction energy, *βJ*. Indeed, for biologically reasonable values of the coupling constant, *βJ* ∼ 0–5, the simulations predict too small a difference between the relaxation times when compared with the experimental data. Fig. 2 shows that for relatively high values of the interaction, *βJ* = 5, cooperativity predicts a relative difference in the relaxation times,

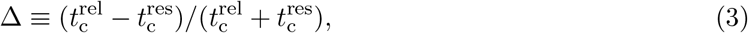

of 20% at most. A simple extrapolation from these results shows that we would need biologically unreasonable values, *βJ >* 19, to obtain values comparable with the experimental ones (∼ 63% for *γ*_500_, and ∼ 82% for *γ*_300_).

Second, taking inspiration from a recent study of depletion effects [10], we constrained the SRLG to interact with a finite reservoir of stator units.We introduced a new parameter, the number of available stator units per cell *n*_tot_, thus fixing the total number of stator units in the system (see SI for more details). The steady-state occupancy increases with *n*_tot_ and tends asymptotically from below to the value of the SRLG model without depletion (Fig. 3). Interestingly, simulations conducted under high depletion (*n*_tot_ = *N*_max_ = 13) predict that release processes relax more rapidly than resurrection ones (Fig. 1), in evident contrast with the experimental results.

**Figure 3:**
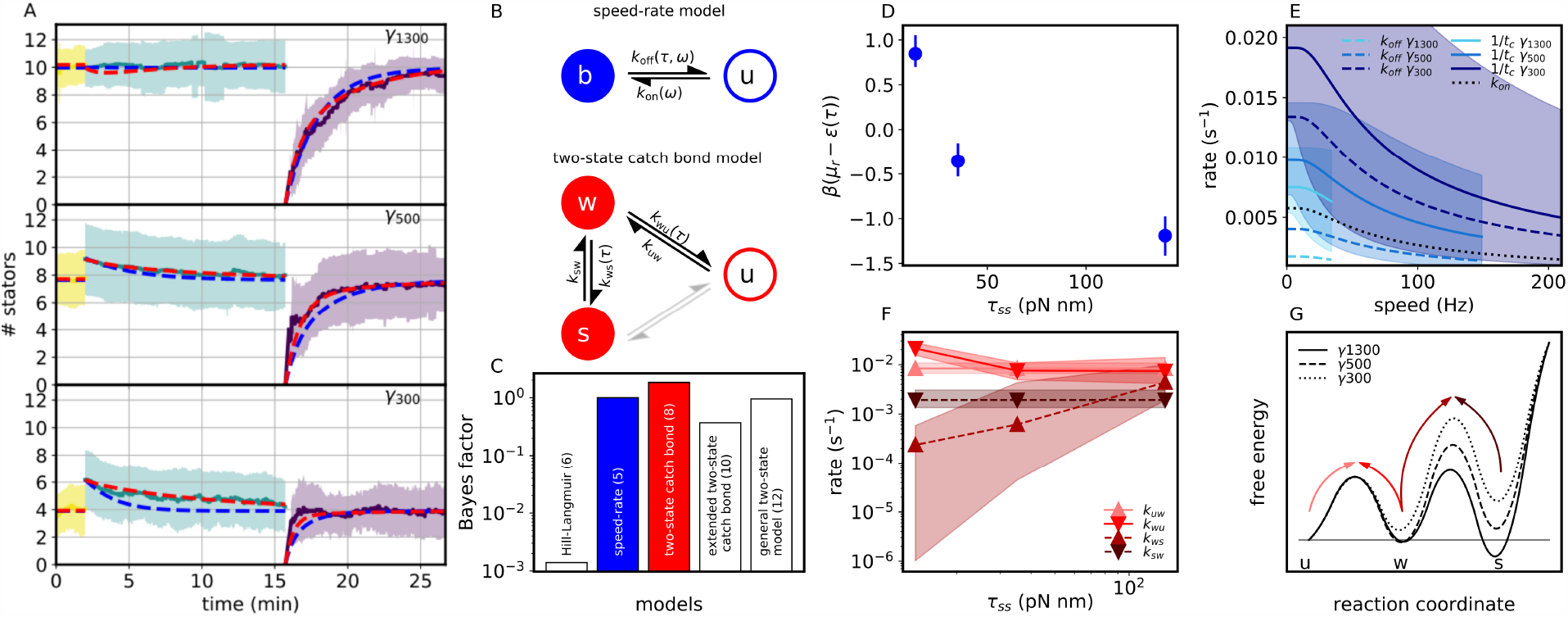
A) The experimental data set. The solid colored lines and shading show the mean and standard deviation of stator number for steady state, release from stall, and resurrection (yellow, green, purple, respectively). Three viscous loads are shown from top to bottom: 1300, 500, and 300 nm bead. Blue dashed lines are a global fit of the speed-rate model of Wadwha et al. [6] (Eq. 1 using the rates of Eq. 5). Red dashed lines are a global fit of the two-state catch-bond model (Eq. 8). B) Schematics of the speed-rate and two-state catch-bond models C) Bayes factors for each of the models, normalized by that of the speed-rate model. Blue and red bars show the speed-rate and two-state catch-bond models depicted in (B). White bars show the 10 and 12 parameter two-state models. D) From the fit of the speed-rate model, the dimensionless energetic contribution *µ*_*r*_ − *ε*(*τ*), as a function of torque per stator unit. Error bars show high density interval. E) From the fit of the speed-rate model, a graphical representation of *k*_on_, *k*_off_ and 1*/t*_*c*_ as a function of motor speed for each of the viscous loads over the range of observed speeds. Shading shows high density interval of 1*/t*_*c*_. F) From the fit of the two-state catch-bond model, rates as a function of torque per stator unit. Triangles mark maximum a priori values, shading shows the high density parameter region. G) From the fit of the two-state catch-bond model, a schematic of the energy landscape corresponding to the rates shown in (F). We note that the distance along the reaction coordinate is unknown. Arrows showing transitions of energy barriers are color-matched to the rates plotted in (F).

In conclusion, all reasonable interpretations of the above findings suggest that neither interactions between neighboring bound stator units, nor finite reservoir effects are directly responsible for the observed asymmetry of relaxation times. Nevertheless, cooperativity and depletion sensibly affect both the steady-state occupancy and the relaxation time. Thus, further theoretical and experimental investigations are needed to ascertain their role in the stator recruitment mechanism.

### A model incorporating speed and torque dependent rates

Wadwha et al. [6] have proposed a model of stator assembly dynamics which employs a statistical physics approach, explicitly incorporating the dependence of the rates on the motor speed, and thus stator stoichiometry (see Fig. 3B). In this model, which we henceforth refer to as the ‘speed-rate’ model, the binding of a single stator unit to the rotor changes its free energy *ε*(*τ*) − *µ*_*r*_, where the binding energy *ε*(*τ*) is dependent upon the torque produced by the stator. In line with our previous model [5], they hypothesize that torque production lowers the free energy difference depending on the torque which leads to the following rate ratio, [6],

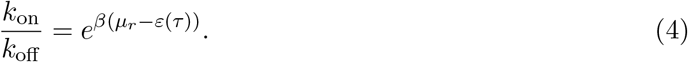

From this general starting point, Wadwha et al. find that their data is best fit by a model in which *k*_off_ is torque dependent, *k*_on_ is speed dependent, and in order to satisfy Eq. 4, *k*_off_ must also incorporate the same speed dependence. The rates are thus

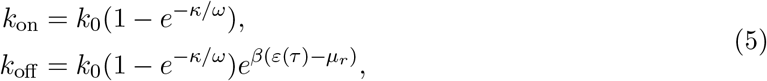

where *k*_0_ and *κ* are constants, and *ω* is the rotation speed of the motor. Because the speed of the motor is proportional to its stoichiometry for a given value of the load (*γ*) [11, 12], the rates of Eq. 5 can also be written in terms of the stoichiometry *κ/ω* ≡ *α*_*γ*_*/N*, where *α*_*γ*_ is a multiplicative constant dependent on the load. This allows us to proceed similarly to the Hill-Langmuir model Eq. 2, and write an ordinary differential equation (ODE) for the evolution of the average stoichiometry,

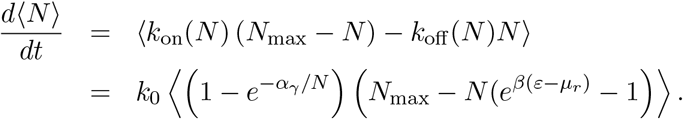

Given the non-linearity of the rates of Eq. 5 with the stoichiometry, the speed-rate model does not give a closed analytical differential equation for the evolution of the average stoichiometry (*N*), as proposed previously [6], since ⟨*f* (*N*)⟩ ≠ *f* (⟨*N*⟩) for any non-linear function *f* (·). As a consequence, calculating the average stoichiometry trajectories for any given set of parameters requires us to computationally solve the corresponding Master Equation (see SI for more details). To infer the optimal parameters of the speed-rate model given the experimental data, we employ Approximate Bayesian Computation (ABC) [13], comparing the resulting average stoichiometry dynamics of the model with the experiments (see SI). The results are shown in Fig. 3A, and the parameters are given in Tab. 3 and Fig. 5. The use of a Bayesian framework not only allows us to perform a global search, reporting the range and correlations of the parameters compatible with the experimental data, but also allows for quantitative comparison of the credibility of various theoretical models (see SI).

### A two-state catch-bond model

We also consider an alternative kinetic model based upon previous models of two-state catch bonds [14–16]. Our previous work observed that the lifetime of the stator in the motor complex increases as the stator applies more torque upon the rotor, and by reaction, as the stator pulls with higher force on the PG [5]. This behavior is characteristic of a catch bond. A widely applied phenomenological model to describe catch-bond behavior is a two-state model in which the thermodynamic stability between two bound conformational states is mediated by force [14, 17]. So, we propose a model, depicted in Fig. 3B, in which the stator can bind to the PG with low affinity or high affinity, mechanical force regulates the transition between these conformational states, and the stator applies torque to the rotor in both bound states. For simplicity, and to minimize the number of free parameters, we assume that the energy barrier between the high affinity (strong, *s*) and unbound (*u*) state is sufficiently high, such that the stator can only transit to and from the unbound state from the low affinity (weak, *w*) state.

The number of weakly bound stators, *w*(*t*), and strongly bound stators, *s*(*t*), follow inherently stochastic dynamics. Similarly to the Hill-Langmuir model (see Eq. 2), we can write the Master Equation for the stochastic process that allows us to obtain a pair of ODEs describing the time evolution of the expected values ⟨*w*⟩ (*t*), and ⟨*s*⟩ (*t*),

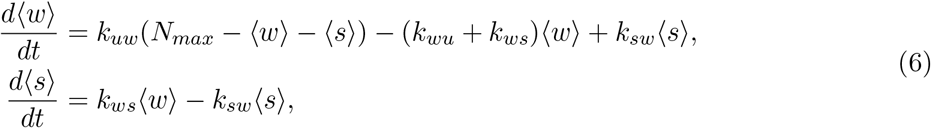

that can be solved to obtain an analytical expression for the stator stoichiometry in time,

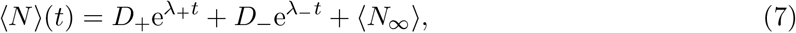

where prefactors *D*_*±*_ depend on the rates and the initial stoichiometry condition, while the steady-state ⟨*N*_*∞*_⟩ and the relaxation rates *λ*_*±*_ only depend on the stator rates *k* (derivation and explicit expressions in SI). Note that, in contrast to the Hill-Langmuir dynamics (Eq. 2), the two-state model naturally predicts two different relaxation time scales in line with the experimental observations.

We hypothesized that the dependence on torque appears in the rates of exiting the intermediate weak state, and we henceforth refer to this as the ‘two-state catch-bond’ model. Nevertheless, in order to generally and more extensively explore two-state models, we also examined cases in which rates exiting the strong state, and binding rates are also allowed to depend upon torque. As in the speed-rate model, we used ABC to infer distributions of parameters compatible with the observed experimental trajectories, with the fit shown in Fig. 3A, parameters listed in Tab. 3, and the posterior parameter distributions shown in Fig. 6-7.

### Model comparison

Despite both dynamical models – the speed-rate model and the two-state catch-bond model – being able to reproduce relaxation asymmetry, the two-state catch-bond model returned average relaxation asymmetries comparable to the experimental ones (Fig. 3A), resulting in trajectories that better fit the data (Fig. 4). This result was anticipated since the two-state catch-bond model involves a larger set of parameters (8) compared to the speed-rate model (5). In order to assess if we can select one model over the other, we calculated the Bayes factor comparing the posterior marginal probabilities of each model by making use of the ABC inferred distributions [18] that naturally incorporate a penalty associated with the dimensionality of the models (see Materials and Methods and SI). Results show that despite having more parameters, the two-state catch-bond model has the highest posterior credibility (see Fig. 3C), though the difference over the speed-rate model is small enough to be considered anecdotal. On the other hand, the difference over other variations of the two-state model (10 and 12 parameter models) is high enough to select the 8 parameter two-state catch-bond model shown in Fig. 3B as the most credible two-state model.

## Discussion

We measured the torque of individual BFMs and calculated the temporal evolution of stator stoichiometry under three sequential conditions: at steady state, after motor stall and subsequent stator recruitment, and during resurrection starting from an empty rotor. These experiments were performed for three viscous loads. We observed that, on average, the relaxation time to steady state is faster for smaller viscous loads, in agreement with our previous work [5]. However, we also observed a subtle but surprising effect that, for each viscous load tested, the relaxation time from ‘below’ (during resurrection) was faster than from ‘above’ (release from stall). This effect is not predicted by our previous simple Hill-Langmuir reversible adsorption model of stator dynamics [5]. We have theoretically explored the effects of interactions between neighboring adsorbed stator units and finite stator unit reservoir effects, and we confirm that neither of these mechanisms can account for the degree of relaxation time asymmetry observed. Such effects may, nonetheless, prove relevant to the description of stator assembly dynamics and are worthy of further exploration (article in preparation [19]).

Based upon previous studies which show that stators dissociate from the motor upon IMF collapse within ∼ one minute, we have assumed that after eight minutes of ionophore-induced PMF collapse, all stator units have dissociated from the rotor and the system is at steady-state zero occupancy [20–22]. This is supported by recent work which estimates the difference in effective free energy of bound and unbound stator units and concludes that the binding of stator units to the motor at zero torque is unfavorable [23].

A recent model of stator dynamics, depicted in Fig. 3B, described by Eqs. 5-6, and referred to here as the ‘speed-rate’ model, proposes that the rate of stator assembly depends non-linearly on both the rotor speed and the torque generated per stator unit [6]. Due to the speed dependency, this model predicts that, when perturbed from steady-state stator stoichiometry, the relaxation time depends upon the direction of perturbation. Our data exhibit this perturbation direction dependent asymmetry in *t*_*c*_ (Fig. 2). Nevertheless, as shown in Figs. 3A and 4, a global fit of this model to our entire data set is still not capable of recapitulating the observed differences in timescales as well as the two-state catch-bond model.

The speed-rate model posits that, when a stator unit binds to the motor, its free energy changes by an amount dependent upon the torque produced by that unit. Fig. 3D shows the resulting change in free energy as a function of single stator torque, with a trend that agrees with Wadwha et al. [6]. Fig. 3E shows the speed-dependent decrease in *k*_on_, *k*_off_, and 1*/t*_*c*_, with the confidence intervals given by ABC. Over the physiological range of speed for *E. coli*, this model would suggest that, as the rotor speeds up, the rate of stator unit binding decreases by about a factor of three. As Wadhwa et al. explain [6], an interaction between the stator unit and the FliG protein of the rotor, triggering a conformational change in the periplasmic region of the stator complex that enables PG binding [24–26], provides a plausible explanation for the rotor speed dependent rate of stator binding. However, the exact mechanism and the specific role of stator-rotor interactions have yet to be clearly illuminated (for a review, see [27]). Importantly, this model also requires that the rate of stator unbinding decreases with increasing motor speed, by the same factor of about three. Speed dependent disassembly is a phenomenon for which we have yet to find a compelling mechanistic explanation.

Our previous work has shown that the stator units have lower rates of unbinding for increased torque upon the rotor, and thus increased force upon the PG, a behavior characteristic of a catch bond [5]. Single molecule force spectroscopy experiments have now identified a wide range of biological catch bonds, particularly amongst proteins with an adhesive or mechanosensory role [28, 29]. While many of the underlying mechanisms which govern this behavior remain to be elucidated, a number of phenomenological and microscopic theories have been proposed [17]. One of the most successful models to date is a two-state model [14, 15] which can quantitatively explain the experimental results for an impressive range of biological catch bonds, including P-selectin [14, 30], the bacterial FimH adhesive protein [16, 31], kinetochore-microtubule interactions [32], cadherin-catenin interactions [33], cell surface sulfatase and glycosaminoglycan interactions [34], vinculin-actin interactions [35], and platelet - von Willebrand factor binding [36].

The two-state catch-bond model (depicted in Fig. 3B) proposes that the stator complex has two bound and torque-producing states, a weak and strong PG affinity state, and that the conversion between these states is force-dependent, with high force favoring the putative high affinity state. Mechanical forces on the stator complex could act directly upon the PG binding domain, for example exposing PG binding sites, in a manner similar to other mechanosensitive proteins [29]. Alternatively, tensile forces could act allosterically via the stalk which links the inner membrane domain to the PG domain. Both experiments and simulations have suggested that allostery plays a role in a number of biological catch bonds, wherein mechanical stress at the allosteric site is propagated along the protein to invoke rearrangements at the binding pocket [29, 37]. We speculate that extension of the disordered and flexible interdomain region of MotB could similarly regulate stator-PG affinity. As seen in Fig. 3A, a global fit of the experimental data to this model produces a very good fit, reproducing the experimentally observed asymmetry in *t*_*c*_. The relaxation time asymmetry would arise because, during resurrection, a stator must merely transition from unbound to weakly bound in order to begin applying torque, whereas during relaxation from stall, stators which are in the strongly bound state must pass through the weakly bound state and then unbind in order to stop producing torque. As depicted in the energy landscape schematic of Fig. 3G (see also Tab. 3 and Figs. 6-8), Bayesian inference in a general two-state model suggests that it is the transition out of the intermediate weak state that is torque-dependent. Increasing torque reduces the barrier between the weak and strong states, in a manner consistent with traditional slip bond behavior. While it is known that the stator must undergo a large conformational change in order to bind to the PG [38, 39], we note that, to date, there is no evidence for two different PG-bound conformations, and this model remains hypothetical. In any case, the success of the two-state model may point towards a more continuous (rather than bistable) or complex conformational and energy landscape for the stator.

Thus, we find that our experimental data fits the two-state catch-bond model more accurately than the speed-rate model. Nevertheless, since the catch-bond model requires a larger dimensional parameter set, we compared the likelihood of both models. While the two-state catch-bond model shows the highest posterior credibility, the difference is insufficient to exclude the speed-rate model.

Neither model offers a specific structural explanation for the catch-bond behavior. The high resolution structures of the stator complex have recently been solved via cryo electron microscopy [40, 41], opening the door to future structure-based models. Simulations may yield atomistic insights regarding the effects of force-induced structural changes on stator assembly. For example, steered molecular dynamics (SMD) simulations, seeded with the static cryo-EM structure, were used to predict force-induced allosteric structural changes in FimH [42], which were later confirmed by single molecule atomic force microscopy experiments [43]. One current limitation to such an approach is that, due to the short timescales accessible, SMD simulations require forces and loading rates much higher than those used in experiments in order to observe bond rupture or allosteric change [17].

In a more recent set of electrorotation experiments, Wadwha et al. revert back to the Hill-Langmuir model [5] to fit their data and find that the extracted rate constants show a speed dependence in *k*_on_ [23], an effect which may be compatible with one of the two models described above. Interestingly, recent electrorotation experiments by Ito et al. also suggest a speed dependence in *k*_on_, albeit in the opposite direction: Ito et al. observe that stator binding is enhanced by rotor speed, though only at low stoichiometry, potentially only applicable to the first stator unit to bind [10]. While we sometimes observe long dwells at zero stoichiometry, potentially compatible with their observations, given our uncertainty in the (short) time needed to restore the PMF during our resurrection experiments, we have not attempted to quantify this effect. We nonetheless hypothesize that this observation, instead of arising from a rotor speed-dependence in *k*_on_, may be equally well explained by the two-state catch-bond model, wherein the transition to the strongly bound state is dependent on the force across the arriving stator unit, which remains at zero until the rotor begins to rotate. We also note that such an effect could arise from the first stator unit needing to recruit a putative partner (for example, FliL [44]), or if the torque from a single stator unit were insufficient to maintain motor rotation [45].

Finally, we note that there is evidence for IMF-dependent stator assembly [46] (as also shown by our resurrection experiments), suggesting that stators sense not only the mechanical environment, but also the electrochemical environment. While this phenomena has yet to be robustly characterized, a successful model of stator assembly dynamics will also need to be able to recapitulate this effect. A catch-bond mechanism within the stator would allow it to stabilize attachment to the PG exactly when it’s needed, provide resistance to large mechanical stresses, and destabilize attachment when it’s no longer needed, allowing reconfiguration under small stresses, thereby conserving the PMF. This mechanism may be consistent with IMF-dependent assembly, though future experiments are needed to quantify the dependence. Future structural models may yield testable predictions linking stator structure and assembly and may even allow the bond to be engineered. Interfering with the catch-bond behavior of the stator could have grand implications for infection, biofilm formation, and disease.

## 1 Materials and Methods

### Bacteria and growth

We used *E. coli* strain MTB32, wherein the flagella has been genetically deleted and the hook is biotinylated [47]. We genetically deleted *cheY* [5], the chemotactic response regulator. Frozen aliquots of cells (grown to saturation and stored in 25% glycerol at -80°C) were grown in Terrific Broth (Sigma-Aldrich) at 33°C for 5 h, shaking at 200 rpm, to a final OD_600_ of 0.5–0.6, then washed and resuspended in motility buffer (10 mM potassium phosphate, 0.1 mM EDTA, 10 mM lactic acid, pH 7.0). Cells were immobilized to a poly-L-lysine (Sigma-Aldrich P4707) coated coverslip in a custom flowslide with a parafilm spacer. Streptavidin superparamagnetic beads (1.36 µm, Sigma-Aldrich; 543 or 302 nm, Adamtech) were washed in PBS, resuspended in motility buffer, then allowed to attach to the BFM hooks.

### BFM experimental measurements

Experiments were performed in motility buffer at 22°C. The sample was illuminated with a 660 nm LED on a custom inverted microscope, and imaged with a 100× 1.45-NA objective (Nikon) onto a CMOS camera (Optronics CL600×2/M) at a framerate of 1 kHz. Two permanent magnets were positioned above the sample at a distance that was controlled by a motorized vertical translation stage.

For a given motor, steady-state rotation was measured with the magnets far from the sample (8 min), the magnets were lowered to within ∼ 1 mm of the sample to stall motor rotation (10 min), the magnets were raised and the motors were allowed to relax to steady state (11 min), the ionophore carbonyl cyanide m-chlorophenyl hydrazone (CCCP, 20 µM) was added to the media to collapse the PMF and dissociate the stators (8 min), then CCCP was washed from the media and cells were allowed to resurrect (11 min). In some measurements of the smaller beads, torque from the magnetic tweezers was insufficient to hold the motor stalled for the entire 10 min. While this effect likely reduces the number of stators recruited during stall, it does not affect the relaxation after stall. In approximately 20% of cases, motors either failed to resurrect after treatment with CCCP or resurrected to less than 40% the steady-state speed; in these cases the entire recording for that motor was discarded.

### Data analysis

The *x, y* position of the bead was determined via image cross-correlation analysis [48], and the angular position of the bead was calculated as *θ* = arctan(*y/x*). The rotational viscous drag coefficient of the bead was calculated as [49],

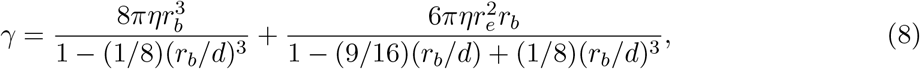

where *r*_*b*_ is the bead radius, *r*_*e*_ is the measured radial distance to the bead’s axis of rotation, and *d* is the distance from the bead to the cell surface, estimated to be 5 nm. Motor torque was calculated as *τ* = *γω*, where *ω* is the rotational speed of the motor. Stator stoichiometry, and single stator torque values were calculated as described previously [5].

All analysis was performed with custom LabView, Matlab, and Python scripts.

### Model simulation

Mean trajectories ⟨*N* (*t, N*_0_)⟩ for an initial motor configuration *N*_0_ for the Hill-Langmuir model and the two-state model were obtained using their analytical expressions (Eqs. 2 and 9). Trajectories for the speed-rate model were obtained by solving the corresponding Master Equation consisting of a system of *N*_max_ linear ordinary differential equations by diagonalizing the resulting rate matrix (see SI).

The initial condition for resurrection trajectories was set to correspond with an empty motor (*N*_0_ = 0), while for stall trajectories the steady-state stoichiometry was used (*N*_0_ = ⟨*N*_*∞*_⟩). In order to reproduce the observed variability of initial conditions during release trajectories, the average trajectory ⟨*N* (*t*)⟩ was obtained by calculating the average over all the initial condition 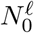 observed for each observed experimental motor *𝓁* = 1, …, *L*,

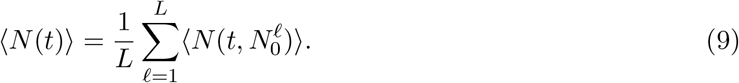

Finally, for the two-state model, release trajectories require inference of the initial values of weakly and strongly bound motors *w*_0_ and *s*_0_ for each experimental observed value of *N*_0_. This relationship was obtained by assuming that the initial configurations in the release experiment is the steady-state probability distribution of occupancy when stator detachment is forbidden *k*_*wu*_ = 0 (see SI).

### Parameter inference and model selection

In order to infer sets of parameters able to reproduce the experimental data, we made use of Approximate Bayesian Computation. The outcome of this analysis returns distributions of credibility for the ensemble of parameters and models with which we obtain intervals and correlations between parameters as well as a means of model comparison. The inference was performed using Sequential Monte Carlo (SMC) to obtain distributions for the credibility *P* (*θ*_*m*_|*d*(*θ*_*m*_) *< ε*) for the parameter set *θ*_*m*_ for each model *m* such that the score *d* of a given parameter set is below a certain threshold *ε* (see SI). The score function *d*(*θ*) was defined as the distance between the experimental trajectories ⟨*N*_exp_(*c, t*)⟩ for a given experimental setup *c* = {resurrection, steady-state, release} and the corresponding predicted trajectories ⟨*N*_theo_(*θ*_*m*_, *t, c*)⟩

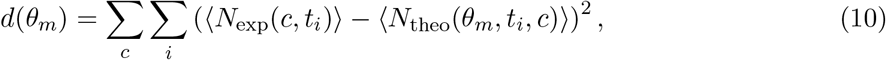

where the subindex *i* runs for all experimental timepoints *t*_*i*_ that include information of at least 3 individual motors with non-zero speed, and a time step of Δ*t* ≡ *t*_*i*+1_ − *t*_*i*_ = 1s. The Bayes factor was calculated by resampling the ABC-SMC [18] with a distance threshold 10% above the minimal threshold for the speed-rate model and using a Gaussian kernel with covariance equal to the the covariance of the original sample. All fittings and SMC simulations were performed with custom Python scripts that can be found in a githhub repository https://github.com/2piruben/BFM_multistate.

## Acknowledgments

We thank Francesco Pedaci for insightful discussions. The bacterial strain used in this work was a modified strain of that gifted to us from the lab of Richard Berry. This work was supported by the ANR FlagMotor project grant ANR-18-CE30-0008 of the French *Agence Nationale de la Recherche*. The CBS is a member of the France-BioImaging (FBI) and the French Infrastructure for Integrated Structural Biology (FRISBI), 2 national infrastructures supported by the French National Research Agency (ANR-10-INBS-04-01 and ANR-10-INBS-05, respectively). AP acknowledges the CNRS for an exemption of a semester (demi-délégation) of teaching duties.

## Notes

### Competing Interest Statement

The authors have declared no competing interest.

