## Supplementary Information for "Relaxation time asymmetry in stator dynamics of the bacterial flagellar motor"

### Stator cooperativity without depletion

We model the rotor as a one-dimensional lattice of $N_{\max}$ sites with periodic boundary conditions in contact with an infinite reservoir of stators described as an ideal solution with constant (dimensionless) chemical potential $\beta\mu_{\mathrm{r}} = \mathrm{cst}$, where $\beta = (k_{\mathrm{B}}T)^{-1}$. Each stator interacts twofold: with the substrate through a (dimensionless) site-independent binding energy $\beta\varepsilon \leq 0$, and, once bound, with its next-neighbor stator through a (dimensionless) interaction energy $\beta J \geq 0$. We associate with each lattice site a variable $\varphi_k$ ($k = 1, \ldots, N_{\max}$) whose value is equal to 0, when the site $k$ is empty, and to +1, when it is occupied by a stator unit. Thus, the total number of adsorbed stator units is $N = \sum_{k=1}^{N_{\max}} \varphi_k$.

This system falls into the universality class of the so-called one-dimensional short range lattice gas (SRLG) models, and its energy depends on the specific configuration $\{\varphi_k\}$ of the lattice sites according to the effective one-dimensional Hamiltonian for bound particles:

$$\mathcal{H}\{\varphi_k\} = -J \sum_{k=1}^{N_{\max}} \varphi_k \varphi_{k+1} - \frac{\mu_{\mathrm{r}}'}{2} \sum_{k=1}^{N_{\max}} (\varphi_{k+1} + \varphi_k) \,. \tag{1}$$

Here, we made use of the periodic boundary conditions, $\varphi_{N_{\max}+1} = \varphi_1$. Furthermore, to simplify the notation, we incorporated the chemical potential of the reservoir and the binding energy into an effective (dimensionless) reservoir chemical potential $\beta\mu_{\mathrm{r}}' = \beta\mu_{\mathrm{r}} - \beta\varepsilon$.

We performed Markov Chain Monte Carlo simulations by applying the Glauber algorithm to the 1D Hamiltonian (Eq. 1) to determine the time evolution of the occupancy expectation value $\langle N(t)\rangle$. After each Monte Carlo step, the average was computed over $10^4$ thermal histories. The initial conditions for resurrection were $\{\varphi_k(0) = 0\}$, implying $N_0 = N(0) = 0$. For the release from stall, we prepared the system such that, for $t < 0$, the statistical ensemble was in equilibrium with its average occupancy matching the desired initial value, thus mimicking the variability of the initial conditions observed in experiments. Then, at $t = 0$, we changed the value of $\beta\mu_{\mathrm{r}}'$, keeping $\beta J$ fixed, to let the system relax towards the same steady state as for the resurrection.

### Stator cooperativity with depletion

Let us consider the case in which the total number of available stators in the system is finite and constant: $n_{\mathrm{tot}} = N(t) + (n_{\mathrm{tot}} - N(t))$. If $N(t)$ denotes the instantaneous number of adsorbed stators

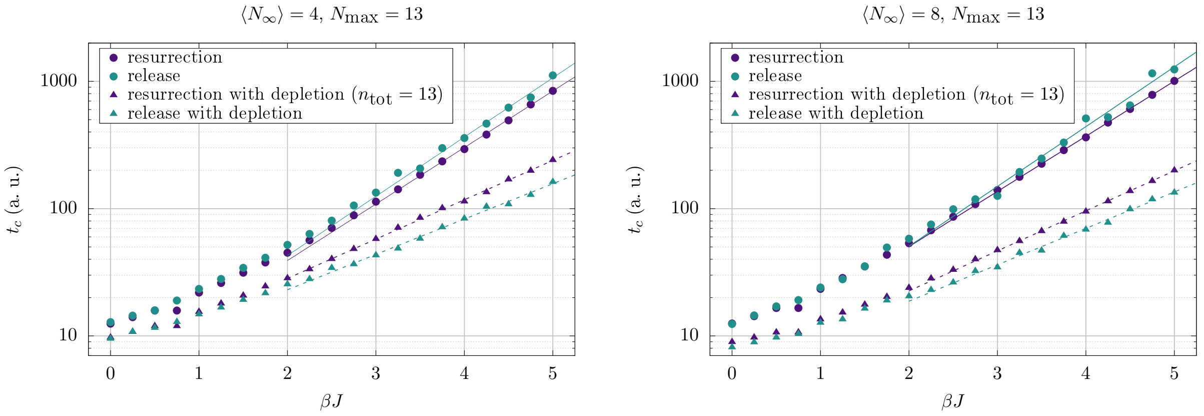


**Figure 1:** Stator cooperativity with (triangles) and without (circles) depletion: relaxation times in arbitrary units (a. u.) versus the coupling constant $\beta J$. $N_{\max} = 13$, and for the depletion case, we have set $n_{\text{tot}} = 13$. The free parameter $\beta\mu'_{\text{r}}$ ($\beta\mu'_0$, in the case with depletion) was adjusted to obtain a steady state occupancy, $\langle N_\infty \rangle$, equal to 4 (left panel) and 8 (right panel), respectively. Green symbols: release from stall, i. e. $\langle N_0 \rangle = 6$ (left) and $\langle N_0 \rangle = 9$ (right panel). Purple symbols: resurrection, i. e. $N_0 = 0$ for both panels. Solid (without depletion) and dashed (with depletion) lines represent the best exponential fit for the data set restricted to values of $J \geq 2$, for which we expect an exponential behavior, as shown theoretically at half-filling [3, 4], i. e. $\langle N_\infty \rangle / N_{\max} = 1/2$: see Tab. 1 for the parameter values. We obtained these results from Monte Carlo simulations using Glauber dynamics for the 1D SRLG model with periodic boundary conditions.

at time $t$, then the instantaneous number of free ones evolves in time too and equals $n_{\text{tot}} - N(t)$. Thus, the chemical potential of the reservoir, $\mu_{\text{r}}$, is no longer constant throughout the relaxation. On the contrary, it decreases as the occupancy increases and *vice versa*. By making the simplifying assumption that the stators behave like a dilute ideal solution, we obtain [1, 2]:

$$\beta\mu_{\text{r}}(t) = \beta\mu_0 + \log\left[n_{\text{tot}} - N(t) + 1\right], \tag{2}$$

where $\mu_0$ corresponds to the chemical potential of the pure solvent. In the case of depletion we can therefore define the effective time-dependent chemical potential of the reservoir as $\beta\mu'_{\text{r}}(t) = \beta\mu_{\text{r}}(t) - \beta\varepsilon$ and consider the difference $\beta\mu'_0 = \beta\mu_0 - \beta\varepsilon$ as an adjustable parameter of the model. To correctly take into account depletion effects, $\beta\mu'_{\text{r}}(t)$ is recalculated after each Monte Carlo step, and the Hamiltonian (Eq. 1) is updated accordingly.

For a given choice of $\beta J$ and $\beta\mu'_0$, the steady state mean occupancy $\langle N_\infty \rangle(\beta J, \beta\mu'_0, n_{\text{tot}})$ increases with the total number of stators in the system, $n_{\text{tot}}$, as one can see in Fig. 3. Depletion strongly affects the evolution of the system for $n_{\text{tot}}/N_{\max} \lesssim 5$. When $n_{\text{tot}}$ increases, depletion effects become less important, and the curves attain the upper bound value corresponding to the steady state mean occupancy without depletion: $\langle N_\infty \rangle(\beta J, \beta\mu'_0, n_{\text{tot}}) \to \langle N_\infty \rangle(\beta J, \beta\mu'_0)$ as $n_{\text{tot}} \to \infty$. Here, we have adjusted $\beta\mu'_0$ such that, for any given $\beta J$, $\langle N_\infty \rangle(\beta J, \beta\mu'_0)$ equals the desired value.

### The Master Equation and average trajectories

All the models described in the manuscript assume that stator binding/unbinding is a stochastic process with rates depending on the occupancy and mechanical properties of the motor. We can accommodate these stochastic descriptions by describing the state of the motor through the probability vector $\vec{P}(t)$, where each component $P^{(i)}(t)$ is the probability of observing the motor at a stoichiometry state $i$ at a time $t$. The different models describe the stator dynamics as a continuous time Markov process in which the probability of observing a change in stoichiometry at a certain time is determined uniquely by the stoichiometry at that time [5]. This entails a linear differential equation for the evolution of $\vec{P}(t)$,

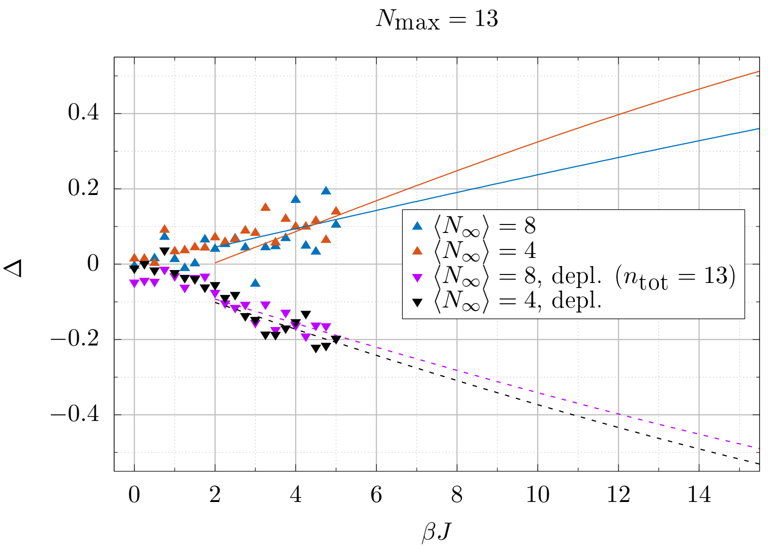


**Figure 2:** Asymmetry in relaxation times, $\Delta \equiv (t_c^{\text{rel}} - t_c^{\text{res}})/(t_c^{\text{rel}} + t_c^{\text{res}})$ (Eq. 3 in the main text): relative difference of relaxation times from release and resurrection processes for mean steady state occupancies $\langle N_\infty \rangle$ of 4 and 8. Same parameter choice and data set as in Fig. 1. Lines represent the predicted relative differences computed by approximating the relaxation times with exponential functions whose parameters are listed in Tab. 1.

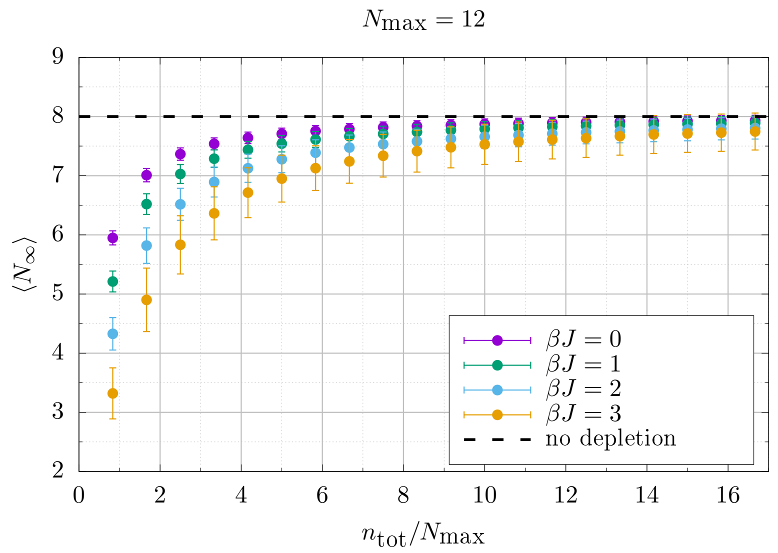


**Figure 3:** Mean steady-state occupancy vs. the total number of stators normalized by $N_{\max}$ for different values of $\beta J$. We choose the free parameter $\beta\mu_0'$ to yield an occupancy $\langle N_\infty \rangle = 2/3 N_{\max} = 8$ (black dashed line) at steady state in the absence of depletion ($n_{\text{tot}} = \infty$). These results were obtained using Monte Carlo simulations with Glauber dynamics of the 1D SRLG model with periodic boundary conditions.

| $N_{\max}$ | $n_{\text{tot}}$ | $\langle N_\infty \rangle$ | Initial conditions | $a \pm \Delta a$ (a. u.) | $b \pm \Delta b$ |
|---|---|---|---|---|---|
| 13 | $\infty$ | 4 | $N_0 = 0$ (res) | $5.1 \pm 0.2$ | $1.022 \pm 0.010$ |
| | | | $\langle N_0 \rangle = 6$ (rel) | $5.0 \pm 0.9$ | $1.07 \pm 0.04$ |
| | | 8 | $N_0 = 0$ (res) | $6.84 \pm 0.14$ | $0.998 \pm 0.004$ |
| | | | $\langle N_0 \rangle = 9$ (rel) | $6 \pm 2$ | $1.08 \pm 0.08$ |
| | 13 | 4 | $N_0 = 0$ (res) | $6.8 \pm 0.3$ | $0.712 \pm 0.009$ |
| | | | $\langle N_0 \rangle = 6$ (rel) | $6.4 \pm 0.5$ | $0.639 \pm 0.019$ |
| | | 8 | $N_0 = 0$ (res) | $5.30 \pm 0.12$ | $0.724 \pm 0.005$ |
| | | | $\langle N_0 \rangle = 9$ (rel) | $5.0 \pm 0.4$ | $0.659 \pm 0.016$ |

**Table 1:** Parameter values $a$ and $b$, with their (asymptotic) standard errors, obtained by best fitting an exponential function, $f(\beta J) = a \exp(\beta J b)$, to the relaxation times $t_c$ from the simulation data (set restricted to data for $J \geq 2$). We denote the absence of depletion by $n_{\text{tot}} = \infty$.

$$\frac{\mathrm{d}\vec{P}}{\mathrm{d}t} = A\vec{P}(t). \tag{3}$$

Where each individual entry $A_{ij}$ of the matrix $A$ describes the linear dependence of a change in the stoichiometry state $i$ on the current probability of occupancy of state $j$.

In the case of the Hill-Langmuir model, the binding propensity is proportional to the available space in the motor $k_{\text{on}}(N_{\max} - N)$, while the detaching propensity is proportional to the current stoichiometry of the motor $k_{\text{off}}N$, where $k_{\text{on}}$ and $k_{\text{off}}$ are the constant binding and unbinding rates, respectively. We can write explicitly the $N_{\max} + 1$ differential equations composing the Master Equation (ME) (Eq. 3) for the Hill-Langmuir model as,

$$\frac{\mathrm{d}P_n}{\mathrm{d}t} = k_{\text{on}}(N_{\max} - n - 1)P_{n-1} + k_{\text{off}}(n+1)P_{n+1} - k_{\text{on}}(N_{\max} - n)P_n - k_{\text{off}} n P_n, \qquad n = 0, \ldots, N_{\max},$$

where $P_{-1} = P_{N_{max}+1} = 0$. The mean occupancy in time $\langle N(t) \rangle = \sum_{n=0}^{N_{\max}} n P_n(t)$ can be obtained by multiplying the above equation by the stoichiometry $n$ and summing for all the possible motor stoichiometries, resulting in a closed differential equation for the evolution of the mean occupancy in time,

$$\frac{d\langle N \rangle}{dt} = k_{\text{on}}\left(N_{\max} - \langle N \rangle\right) - k_{\text{off}} \langle N \rangle. \tag{4}$$

Similarly, we can write the Master Equation for the speed-rate model by introducing the stoichiometry dependent rates $k_{\text{on}}(N)$ and $k_{\text{off}}(N)$ in Eq. 4. If we try to obtain an equation for $\langle N \rangle$ by multiplying the ME by $n$ and summing for all the possible occupancy states of the motor, we do not recover a closed expression for $\langle N \rangle$ due to the non-linearity of the rates with the occupancy,

$$\begin{aligned} \frac{d\langle N \rangle}{dt} &= \langle k_{\text{on}}(N)\left(N_{\max} - N\right) - k_{\text{off}}(N) N \rangle \\ &= k_0 \left\langle \left(1 - e^{-\alpha_\gamma / N}\right)\left(N_{\max} - N(e^{\beta(\varepsilon - \mu_r)} - 1\right)\right\rangle. \end{aligned}$$

Therefore, analysis of the mean occupancy in time demands that we solve the systems of differential equations numerically. Note that, though the rates are not linear in the occupancy $N$, the right hand side of the ODE system is still linear in the probability vector $\vec{P}$ (see eq. 3), and this allows us to solve the equation by diagonalization of the matrix $A$ (see next section).

Finally, for the two-state model, the stoichiometry probability vector must incorporate the two possible binding states (weak and strong) of the stators $\vec{P} = \{P_{w,s} | w = 0, \ldots, N_{\max};\ s = 0, \ldots, N_{\max};\ w + s \leq N_{\max}\}$, where the resulting system of $(N_{\max} - 2)(N_{\max} - 1)/2$ differential equations composing the Master Equation follows,

$$\begin{aligned}
\frac{\mathrm{d}P_{w,s}}{\mathrm{d}t} &= k_{uw}[(N_{\max} - w - s - 1)P_{w-1,s} - (N_{\max} - w - s)P_{w,s}], \\
&+ k_{wu}((w + 1)P_{w+1,s} - wP_{w,s}), \\
&+ k_{ws}((w + 1)P_{w+1,s-1} - wP_{w,s}), \\
&+ k_{sw}((s + 1)P_{w-1,s+1} - sP_{w,s}), \quad w, s = 0, 1, ..., N_{\max},
\end{aligned} \tag{5}$$

where for sake of clarity we are extending the notation such that $P_{w,s} = 0$ if $w, s < 0$ or $w+s > N_{\max}$. Despite the high dimensionality of the two-state model, since the right hand side is linear in the stoichiometry variables $w, s$, we can calculate closed expressions for the evolution of the expected number of weakly bound stators at time $t$ as $\langle w(t) \rangle = \sum_{w=0}^{N_{\max}} \sum_{s=0}^{N_{\max}} wP_{w,s}$; and equivalently, for the strongly bound stators $\langle s(t) \rangle = \sum_{w=0}^{N_{\max}} \sum_{s=0}^{N_{\max}} sP_{w,s}$. Multiplying Eq. 5 by $w$ or by $s$ and summing for all the possible attainable configurations of the motor, we obtain the system of equations

$$\begin{aligned}
\frac{\mathrm{d}\langle w \rangle}{\mathrm{d}t} &= k_{uw}(N_{\max} - \langle w \rangle - \langle s \rangle) + k_{sw}\langle s \rangle - (k_{wu} + k_{ws})\langle w \rangle, \\
\frac{\mathrm{d}\langle s \rangle}{\mathrm{d}t} &= k_{ws}\langle w \rangle - k_{sw}\langle s \rangle.
\end{aligned} \tag{6}$$

At steady state ($\frac{\mathrm{d}\langle w \rangle}{\mathrm{d}t} = \frac{\mathrm{d}\langle w \rangle}{\mathrm{d}t} = 0$), the model predicts the following stoichiometries,

$$\begin{aligned}
\langle w_\infty \rangle &= \frac{N_{\max} k_{sw} k_{uw}}{k_{uw}k_{ws} + k_{uw}k_{sw} + k_{sw}k_{wu}}, \\
\langle s_\infty \rangle &= \frac{N_{\max} k_{ws} k_{uw}}{k_{uw}k_{ws} + k_{uw}k_{sw} + k_{sw}k_{wu}}, \\
\langle N_\infty \rangle = \langle w_\infty + s_\infty \rangle &= \frac{N_{\max}(k_{ws} + k_{sw})k_{uw}}{k_{uw}k_{ws} + k_{uw}k_{sw} + k_{sw}k_{wu}}.
\end{aligned} \tag{7}$$

In addition, the linear system of ODEs (6) can be solved analytically, obtaining the evolution of the weak and strongly bound stators,

$$\begin{pmatrix} \langle w \rangle(t) \\ \langle s \rangle(t) \end{pmatrix} = C_+ \begin{pmatrix} 1 \\ a_+ \end{pmatrix} \mathrm{e}^{\lambda_+ t} + C_- \begin{pmatrix} 1 \\ a_- \end{pmatrix} \mathrm{e}^{\lambda_- t} + \begin{pmatrix} \langle w \rangle_\infty \\ \langle s \rangle_\infty \end{pmatrix}, \tag{8}$$

where,

$$\begin{aligned}
\lambda_\pm &= -\frac{K}{2}\left(1 \mp \sqrt{1 - 4\frac{k_{uw}k_{ws} + k_{uw}k_{sw} + k_{sw}k_{wu}}{K^2}}\right), \\
a_\pm &= \frac{\lambda_\pm + k_{uw} + k_{wu} + k_{ws}}{k_{sw} - k_{uw}} = \frac{\lambda_\mp + k_{sw}}{k_{uw} - k_{sw}}, \\
K &= k_{uw} + k_{wu} + k_{ws} + k_{sw}, \\
C_- &= \frac{(k_{sw} - k_{uw})(s_0 - \langle s \rangle_{ss} - a_+(w_0 - \langle w \rangle_{ss}))}{\lambda_- - \lambda_+}, \\
C_+ &= w_0 - \langle w \rangle_{ss} - C_-.
\end{aligned} \tag{9}$$

The expected trajectory for a given set of rates and initial condition can be obtained from Eq. (8), returning the expression given in the main text,

$$\langle N(t)\rangle = \langle w(t)\rangle + \langle s(t)\rangle = C_+(1+a_+)\mathrm{e}^{\lambda_+ t} + C_-(1+a_-)\mathrm{e}^{\lambda_- t} + \langle w_\infty\rangle + \langle s_\infty\rangle \qquad (10)$$

$$\equiv D_+\mathrm{e}^{\lambda_+ t} + D_-\mathrm{e}^{\lambda_- t} + \langle N_\infty\rangle \qquad (11)$$

### Integration of the Master Equation

Since all the model comparisons in this study are done through the expected occupancy trajectories, the analytical expressions for $\langle N(t)\rangle$ were used when available, without the necessity to solve the Master Equation (Eq. 3). For the speed-rate model this is not possible. Nevertheless, since the Master Equation is linear in the probabilities $P_n$, it can be solved by diagonalizing the system of ODEs finding the change of base $\Lambda = UAU^{-1}$ that returns a diagonal matrix $\Lambda$ decoupling the system of ODEs (where the matrix $U$ is composed by the eigenvectors of $A$). In the eigenbase, the evolution for each component of the probability vector follows,

$$\frac{\mathrm{d}Q_n(t)}{\mathrm{d}t} = \lambda_n Q_n(t) \quad \Rightarrow \quad Q_n(t) = Q_n(0)\mathrm{e}^{\lambda_n t}, \qquad (12)$$

where $\vec{Q}(t) = U\vec{P}(t)$ is the probability vector in the eigenbase and $\lambda_n = \Lambda_{n,n}$ are the corresponding eigenvalues. Full trajectories $\vec{P}(t)$ were obtained by numerically diagonalizing the matrix $A$ for a given parameter set, expressing the initial vector $\vec{P}(0)$ in the eigenbase $\vec{Q}(0) = U\vec{P}(0)$, calculating the trajectories in the eigenbase $\vec{Q}(t)$ using Eq. 12, and finally, transforming back to the natural base $\vec{P}(t) = U^{-1}\vec{Q}(t)$.

This evaluation was required to obtain the trajectories of the speed-rate model. In addition, it was also used to obtain the initial condition for the release trajectories of the two-state model. Experimental release trajectories provide the initial condition $\langle N(0)\rangle = N_{\mathrm{exp}}(0)$ but leave a degree of freedom on the distribution of the stator hidden states $\langle w(0)\rangle$ and $\langle s(0)\rangle = \langle N(0)\rangle - \langle w(0)\rangle$. Different initial conditions were tested, where the best fits were obtained by using the steady state distribution $P^*_{w,s}$ (eigenvector with $\lambda = 0$) in the case for which there is no unbinding from the motor $k_{wu} = 0$. This is consistent with the catch-bond hypothesis, assuming that unbinding is forbidden during stall. The corresponding values $\langle w(0)\rangle$ and $\langle s(0)\rangle = N(0) - \langle w(0)\rangle$ were obtained using the conditional average,

$$\langle w(0)\rangle = \frac{\sum_{w,s=1,\ldots,N_{max}} wP^*_{w,s}\delta_{w+s,N_0}}{\sum_{w,s=1,\ldots,N_{max}} P^*_{w,s}\delta_{w+s,N_0}}, \qquad (13)$$

where $\delta_{i,j}$ is the Kronecker delta.

### Bayesian Inference

Bayesian inference was employed to obtain credibility distributions for each set of parameters for each model. Bayesian inference requires us to define a likelihood function that describes the probability to observe the experimental data given a certain set of parameters. Unfortunately, variability between motors and uncertainty from the stoichiometries obtained by the step detection and stoichiometry reconstruction alogorithms [6] do not allow us to write an exact expression for the likelihood. For this reason, we employed Approximate Bayesian Computation (ABC) in which the likelihood function for a given parameter set $\vec{\theta}$ is replaced by the conditional probability $P(d(\vec{\theta}, data) < \epsilon)$. Where the function $d(\vec{\theta}, data)$ is a distance function comparing the experimental

| Model | Optimal Parameters<br>Optimal distance | <br>Maximum a Posteriori | <br>High density interval |
|---|---|---|---|
| Hill Langmuir<br>release data<br>$u \underset{k_{\text{off}}(\tau)}{\overset{k_{\text{on}}(\tau)}{\rightleftharpoons}} b$<br>$\epsilon_T = 45$<br>$d^* = 35.5$ | $k_{\text{on},300} = 2.18 \cdot 10^{-3}$ s$^{-1}$<br>$k_{\text{off},300} = 3.96 \cdot 10^{-3}$ s$^{-1}$<br>$k_{\text{on},500} = 2.47 \cdot 10^{-3}$ s$^{-1}$<br>$k_{\text{off},500} = 1.57 \cdot 10^{-3}$ s$^{-1}$<br>$k_{\text{on},1300}/k_{\text{off},1300} = 3.54$ | $k_{\text{on},300} = 2.04 \cdot 10^{-3}$ s$^{-1}$<br>$k_{\text{off},300} = 3.73 \cdot 10^{-3}$ s$^{-1}$<br>$k_{\text{on},500} = 2.45 \cdot 10^{-3}$ s$^{-1}$<br>$k_{\text{off},500} = 1.57 \cdot 10^{-3}$ s$^{-1}$<br>$k_{\text{on},1300}/k_{\text{off},1300} = 3.64$ | $k_{\text{on},300} = [1.28, 3.01] \cdot 10^{-3}$ s$^{-1}$<br>$k_{\text{off},300} = [2.47, 5.33] \cdot 10^{-3}$ s$^{-1}$<br>$k_{\text{on},500} = [1.30, 4.78] \cdot 10^{-3}$ s$^{-1}$<br>$k_{\text{off},500} = [9.05, 28.6] \cdot 10^{-4}$ s$^{-1}$<br>$k_{\text{on},1300}/k_{\text{off},1300} = [3.38, 4.02]$ |
| Hill Langmuir<br>resurrection data<br>$\epsilon_T = 190$<br>$d^* = 169$ | $k_{\text{on},300} = 0.0189$ s$^{-1}$<br>$k_{\text{off},300} = 0.0451$ s$^{-1}$<br>$k_{\text{on},500} = 9.89 \cdot 10^{-3}$ s$^{-1}$<br>$k_{\text{off},500} = 8.08 \cdot 10^{-3}$ s$^{-1}$<br>$k_{\text{on},1300} = 5.39 \cdot 10^{-3}$ s$^{-1}$<br>$k_{\text{off},1300} = 1.97 \cdot 10^{-3}$ s$^{-1}$ | $k_{\text{on},300} = 0.0216$ s$^{-1}$<br>$k_{\text{off},300} = 0.0523$ s$^{-1}$<br>$k_{\text{on},500} = 9.62 \cdot 10^{-3}$ s$^{-1}$<br>$k_{\text{off},500} = 7.80 \cdot 10^{-3}$ s$^{-1}$<br>$k_{\text{on},1300} = 5.38 \cdot 10^{-3}$ s$^{-1}$<br>$k_{\text{off},1300} = 1.94 \cdot 10^{-3}$ s$^{-1}$ | $k_{\text{on},300} = [0.0106, 0.0427]$ s$^{-1}$<br>$k_{\text{off},300} = [0.0251, 0.100]$ s$^{-1}$<br>$k_{\text{on},500} = [8.23, 11.7] \cdot 10^{-3}$ s$^{-1}$<br>$k_{\text{off},500} = [6.63, 9.86] \cdot 10^{-3}$ s$^{-1}$<br>$k_{\text{on},1300} = [4.90, 5.87] \cdot 10^{-3}$ s$^{-1}$<br>$k_{\text{off},1300} = [1.61, 2.29] \cdot 10^{-3}$ s$^{-1}$ |
| Hill Langmuir<br>full dataset<br>$\epsilon_T = 1100$<br>$d^* = 1024$ | $k_{\text{on},300} = 6.81 \cdot 10^{-3}$ s$^{-1}$<br>$k_{\text{off},300} = 1.37 \cdot 10^{-2}$ s$^{-1}$<br>$k_{\text{on},500} = 7.08 \cdot 10^{-3}$ s$^{-1}$<br>$k_{\text{off},500} = 4.68 \cdot 10^{-3}$ s$^{-1}$<br>$k_{\text{on},1300} = 4.81 \cdot 10^{-3}$ s$^{-1}$<br>$k_{\text{off},1300} = 1.39 \cdot 10^{-3}$ s$^{-1}$ | $k_{\text{on},300} = 9.21 \cdot 10^{-3}$ s$^{-1}$<br>$k_{\text{off},300} = 1.86 \cdot 10^{-2}$ s$^{-1}$<br>$k_{\text{on},500} = 7.11 \cdot 10^{-3}$ s$^{-1}$<br>$k_{\text{off},500} = 4.74 \cdot 10^{-3}$ s$^{-1}$<br>$k_{\text{on},1300} = 4.77 \cdot 10^{-3}$ s$^{-1}$<br>$k_{\text{off},1300} = 1.38 \cdot 10^{-3}$ s$^{-1}$ | $k_{\text{on},300} = [4.36, 16.8] \cdot 10^{-3}$ s$^{-1}$<br>$k_{\text{off},300} = [8.82, 33.5] \cdot 10^{-3}$ s$^{-1}$<br>$k_{\text{on},500} = [5.69, 9.21] \cdot 10^{-3}$ s$^{-1}$<br>$k_{\text{off},500} = [3.71, 6.07] \cdot 10^{-3}$ s$^{-1}$<br>$k_{\text{on},1300} = [4.26, 5.45] \cdot 10^{-3}$ s$^{-1}$<br>$k_{\text{off},1300} = [1.11, 1.65] \cdot 10^{-3}$ s$^{-1}$ |

**Table 2:** Resulting parameters for the Hill-Langmuir model [6]. The SMC-ABC was run for each case with a final distance threshold $\epsilon_T$ and the optimal distance parameter set returned an optimal distance $d^*$.

average trajectories, with the expected average trajectories for $\vec{\theta}$. We used the sum of squared distances between experimental and theoretical average trajectories (see Material and Methods). The threshold $\epsilon$ is a small value below which a trajectory is considered to reproduce experimental trajectories.

The inference also requires the introduction of a prior knowledge of the model expressed as prior credibility distributions for each parameter. To avoid bias in the parameter inference, we chose flat distributions for the logarithm of each rate over three decades. To make this clearer, the range of all posterior plots (see Figs. 5-8) shows the range of the prior distributions used.

In order to accelerate the convergence of the inference we made use of Sequential Monte Carlo (SMC) sampling in which the posterior distribution is obtained by sequentially finding the posterior distributions for a decreasing set of thresholds $\epsilon_1, \epsilon_2, \ldots, \epsilon_T$ obtaining a set of generations of posteriors until the target threshold $\epsilon_T$ is obtained. Perturbation and evaluation for each generation was done using a Gaussian Kernel with a covariance dependent on the previous generation [7].

For each individual model, the final threshold value $\epsilon_T$ was chosen to be close to the optimal distance attainable for each model. On the other hand, for model selection, the SMC-ABC was run with a common final threshold 10% above the optimal distance for the speed-rate model.

### Fitting results of the experimental data

For each posterior distribution, we report in tables 2 and 3 the maximum a posteriori of the distribution and the 90% high density intervals for each parameter set. In addition, for each parameter set, we performed a minimization of the distance function, using the Nedler-Mead algorithm, starting each search at the maximum a posteriori identified by the SMC-ABC.

| Model | Optimal Parameters<br>Optimal distance | <br>Maximum a Posteriori | <br>High density interval |
|---|---|---|---|
| speed-rate [8]<br>(5 parameters)<br>$\mathrm{u} \underset{k_{\mathrm{off}}(\omega,\tau)}{\overset{k_{\mathrm{on}}(\omega)}{\rightleftharpoons}} \mathrm{b}$<br>$\epsilon_T = 1100$<br>$d^* = 997$ | $k_0 = 0.00575\ \mathrm{s}^{-1}$<br>$\kappa = 396\ \mathrm{rad\ s}^{-1}$<br>$\beta(\mu_r - \varepsilon_{300}) = 0.846$<br>$\beta(\mu_r - \varepsilon_{500}) = -0.356$<br>$\beta(\mu_r - \varepsilon_{1300}) = -1.19$ | $k_0 = 0.00562\ \mathrm{s}^{-1}$<br>$\kappa = 256\ \mathrm{rad\ s}^{-1}$<br>$\beta(\mu_r - \varepsilon_{300}) = 0.871$<br>$\beta(\mu_r - \varepsilon_{500}) = -0.336$<br>$\beta(\mu_r - \varepsilon_{1300}) = -1.20$ | $k_0 = [0.00430, 0.0079]\ \mathrm{s}^{-1}$<br>$\kappa = [109, 818]\ \mathrm{rad\ s}^{-1}$<br>$\beta(\mu_r - \varepsilon_{300}) = [0.712, 1.04]$<br>$\beta(\mu_r - \varepsilon_{500}) = [-0.516, -0.168]$<br>$\beta(\mu_r - \varepsilon_{1300}) = [-1.40, -0.990]$ |
| two-state catch-bond<br>(8 parameters)<br>$\mathrm{u} \underset{k_{wu}(\tau)}{\overset{k_{uw}}{\rightleftharpoons}} \mathrm{w} \underset{k_{sw}}{\overset{k_{ws}(\tau)}{\rightleftharpoons}} \mathrm{s}$<br>$\epsilon_T = 500$<br>$d^* = 391$ | $k_{uw} = 8.37 \cdot 10^{-3}\ \mathrm{s}^{-1}$<br>$k_{sw} = 1.93 \cdot 10^{-3}\ \mathrm{s}^{-1}$<br>$k_{wu,300} = 0.0213\ \mathrm{s}^{-1}$<br>$k_{ws,300} = 2.37 \cdot 10^{-4}\ \mathrm{s}^{-1}$<br>$k_{wu,500} = 7.53 \cdot 10^{-3}\ \mathrm{s}^{-1}$<br>$k_{ws,500} = 6.17 \cdot 10^{-4}\ \mathrm{s}^{-1}$<br>$k_{wu,1300} = 7.42 \cdot 10^{-3}\ \mathrm{s}^{-1}$<br>$k_{ws,1300} = 4.38 \cdot 10^{-3}\ \mathrm{s}^{-1}$ | $k_{uw} = 8.03 \cdot 10^{-3}\ \mathrm{s}^{-1}$<br>$k_{sw} = 2.34 \cdot 10^{-3}\ \mathrm{s}^{-1}$<br>$k_{wu,300} = 0.0194\ \mathrm{s}^{-1}$<br>$k_{ws,300} = 3.63 \cdot 10^{-5}\ \mathrm{s}^{-1}$<br>$k_{wu,500} = 6.90 \cdot 10^{-3}\ \mathrm{s}^{-1}$<br>$k_{ws,500} = 4.23 \cdot 10^{-4}\ \mathrm{s}^{-1}$<br>$k_{wu,1300} = 7.94 \cdot 10^{-3}\ \mathrm{s}^{-1}$<br>$k_{ws,1300} = 6.02 \cdot 10^{-3}\ \mathrm{s}^{-1}$ | $k_{uw} = [6.67, 10.8] \cdot 10^{-3}\ \mathrm{s}^{-1}$<br>$k_{sw} = [1.35, 3.06] \cdot 10^{-3}\ \mathrm{s}^{-1}$<br>$k_{wu,300} = [1.54, 2.74] \cdot 10^{-2}\ \mathrm{s}^{-1}$<br>$k_{ws,300} = [1.02, 589] \cdot 10^{-6}\ \mathrm{s}^{-1}$<br>$k_{wu,500} = [4.93, 10.9] \cdot 10^{-3}\ \mathrm{s}^{-1}$<br>$k_{ws,500} = [4.46, 134] \cdot 10^{-5}\ \mathrm{s}^{-1}$<br>$k_{wu,1300} = [4.13, 13.9] \cdot 10^{-3}\ \mathrm{s}^{-1}$<br>$k_{ws,1300} = [1.91, 9.81] \cdot 10^{-3}\ \mathrm{s}^{-1}$ |
| extended two-state catch-bond<br>(10 parameters)<br>$\mathrm{u} \underset{k_{wu}(\tau)}{\overset{k_{uw}}{\rightleftharpoons}} \mathrm{w} \underset{k_{sw}(\tau)}{\overset{k_{ws}(\tau)}{\rightleftharpoons}} \mathrm{s}$<br>$\epsilon_T = 500$<br>$d^* = 388$ | $k_{uw} = 8.41 \cdot 10^{-3}\ \mathrm{s}^{-1}$<br>$k_{wu,300} = 0.0212\ \mathrm{s}^{-1}$<br>$k_{ws,300} = 2.49 \cdot 10^{-4}\ \mathrm{s}^{-1}$<br>$k_{sw,300} = 2.12 \cdot 10^{-3}\ \mathrm{s}^{-1}$<br>$k_{wu,500} = 7.55 \cdot 10^{-3}\ \mathrm{s}^{-1}$<br>$k_{ws,500} = 5.90 \cdot 10^{-4}\ \mathrm{s}^{-1}$<br>$k_{sw,500} = 1.87 \cdot 10^{-3}\ \mathrm{s}^{-1}$<br>$k_{wu,1300} = 7.23 \cdot 10^{-3}\ \mathrm{s}^{-1}$<br>$k_{ws,1300} = 3.88 \cdot 10^{-3}\ \mathrm{s}^{-1}$<br>$k_{sw,1300} = 1.76 \cdot 10^{-3}\ \mathrm{s}^{-1}$ | $k_{uw} = 8.00 \cdot 10^{-3}\ \mathrm{s}^{-1}$<br>$k_{wu,300} = 0.0192\ \mathrm{s}^{-1}$<br>$k_{ws,300} = 6.45 \cdot 10^{-5}\ \mathrm{s}^{-1}$<br>$k_{sw,300} = 2.01 \cdot 10^{-3}\ \mathrm{s}^{-1}$<br>$k_{wu,500} = 7.04 \cdot 10^{-3}\ \mathrm{s}^{-1}$<br>$k_{ws,500} = 4.18 \cdot 10^{-4}\ \mathrm{s}^{-1}$<br>$k_{sw,500} = 1.69 \cdot 10^{-3}\ \mathrm{s}^{-1}$<br>$k_{wu,1300} = 6.07 \cdot 10^{-3}\ \mathrm{s}^{-1}$<br>$k_{ws,1300} = 2.84 \cdot 10^{-3}\ \mathrm{s}^{-1}$<br>$k_{sw,1300} = 1.66 \cdot 10^{-3}\ \mathrm{s}^{-1}$ | $k_{uw} = [6.54, 10.5] \cdot 10^{-3}\ \mathrm{s}^{-1}$<br>$k_{wu,300} = [0.0152, 0.0254]\ \mathrm{s}^{-1}$<br>$k_{ws,300} = [1.06, 666.0] \cdot 10^{-6}\ \mathrm{s}^{-1}$<br>$k_{sw,300} = [1.30, 4.21] \cdot 10^{-3}\ \mathrm{s}^{-1}$<br>$k_{wu,500} = [4.80, 10.2] \cdot 10^{-3}\ \mathrm{s}^{-1}$<br>$k_{ws,500} = [6.27, 163] \cdot 10^{-5}\ \mathrm{s}^{-1}$<br>$k_{sw,500} = [7.72, 42.8] \cdot 10^{-4}\ \mathrm{s}^{-1}$<br>$k_{wu,1300} = [3.72, 12.1] \cdot 10^{-3}\ \mathrm{s}^{-1}$<br>$k_{ws,1300} = [1.15, 9.77] \cdot 10^{-3}\ \mathrm{s}^{-1}$<br>$k_{sw,1300} = [8.24, 34.3] \cdot 10^{-4}\ \mathrm{s}^{-1}$ |
| general two-state model<br>(12 parameters)<br>$\mathrm{u} \underset{k_{wu}(\tau)}{\overset{k_{uw}(\tau)}{\rightleftharpoons}} \mathrm{w} \underset{k_{sw}(\tau)}{\overset{k_{ws}(\tau)}{\rightleftharpoons}} \mathrm{s}$<br>$\epsilon_T = 325$<br>$d^* = 222$ | $k_{uw,300} = 0.0697\ \mathrm{s}^{-1}$<br>$k_{wu,300} = 0.188\ \mathrm{s}^{-1}$<br>$k_{ws,300} = 4.76 \cdot 10^{-4}\ \mathrm{s}^{-1}$<br>$k_{sw,300} = 2.06 \cdot 10^{-3}\ \mathrm{s}^{-1}$<br>$k_{uw,500} = 0.0127 \cdot\ \mathrm{s}^{-1}$<br>$k_{wu,500} = 0.0132\ \mathrm{s}^{-1}$<br>$k_{ws,500} = 1.02 \cdot 10^{-3}\ \mathrm{s}^{-1}$<br>$k_{sw,500} = 1.82 \cdot 10^{-3}\ \mathrm{s}^{-1}$<br>$k_{uw,1300} = 6.15 \cdot 10^{-3}\ \mathrm{s}^{-1}$<br>$k_{wu,1300} = 4.07 \cdot 10^{-3}\ \mathrm{s}^{-1}$<br>$k_{ws,1300} = 3.61 \cdot 10^{-3}\ \mathrm{s}^{-1}$<br>$k_{sw,1300} = 2.62 \cdot 10^{-3}\ \mathrm{s}^{-1}$ | $k_{uw,300} = 0.0252\ \mathrm{s}^{-1}$<br>$k_{wu,300} = 0.0589\ \mathrm{s}^{-1}$<br>$k_{ws,300} = 2.76 \cdot 10^{-5}\ \mathrm{s}^{-1}$<br>$k_{sw,300} = 2.21 \cdot 10^{-3}\ \mathrm{s}^{-1}$<br>$k_{uw,500} = 0.0124\ \mathrm{s}^{-1}$<br>$k_{wu,500} = 0.0130\ \mathrm{s}^{-1}$<br>$k_{ws,500} = 7.60 \cdot 10^{-4}\ \mathrm{s}^{-1}$<br>$k_{sw,500} = 1.33 \cdot 10^{-3}\ \mathrm{s}^{-1}$<br>$k_{uw,1300} = 5.51 \cdot 10^{-3}\ \mathrm{s}^{-1}$<br>$k_{wu,1300} = 2.75 \cdot 10^{-3}\ \mathrm{s}^{-1}$<br>$k_{ws,1300} = 2.14 \cdot 10^{-3}\ \mathrm{s}^{-1}$<br>$k_{sw,1300} = 2.75 \cdot 10^{-3}\ \mathrm{s}^{-1}$ | $k_{uw,300} = [0.00906, 0.0978]\ \mathrm{s}^{-1}$<br>$k_{wu,300} = [0.0209, 0.234]\ \mathrm{s}^{-1}$<br>$k_{ws,300} = [1.00, 434] \cdot 10^{-6}\ \mathrm{s}^{-1}$<br>$k_{sw,300} = [1.09, 3.51] \cdot 10^{-3}\ \mathrm{s}^{-1}$<br>$k_{uw,500} = [7.94, 19.9] \cdot 10^{-3}\ \mathrm{s}^{-1}$<br>$k_{wu,500} = [6.74, 23.5] \cdot 10^{-3}\ \mathrm{s}^{-1}$<br>$k_{ws,500} = [2.18, 29] \cdot 10^{-4}\ \mathrm{s}^{-1}$<br>$k_{sw,500} = [7.81, 35.7] \cdot 10^{-4}\ \mathrm{s}^{-1}$<br>$k_{uw,1300} = [4.67, 7.58] \cdot 10^{-3}\ \mathrm{s}^{-1}$<br>$k_{wu,1300} = [1.83, 6.80] \cdot 10^{-3}\ \mathrm{s}^{-1}$<br>$k_{ws,1300} = [3.57, 94.1] \cdot 10^{-4}\ \mathrm{s}^{-1}$<br>$k_{sw,1300} = [4.29, 98.4] \cdot 10^{-4}\ \mathrm{s}^{-1}$ |

**Table 3:** Resulting parameters for the two different models which show asymmetric relaxation timescales discussed in the manuscript: the speed-rate model, and the two-state catch bond model. In addition we include the results for the extended two-state catch bond model (10 parameters) where only the binding rate is independent of the torque, and the general two-state model (12 parameters). The SMC-ABC was run with a final distance threshold $\epsilon_T$ and the optimal distance parameter set returned an optimal distance $d^*$

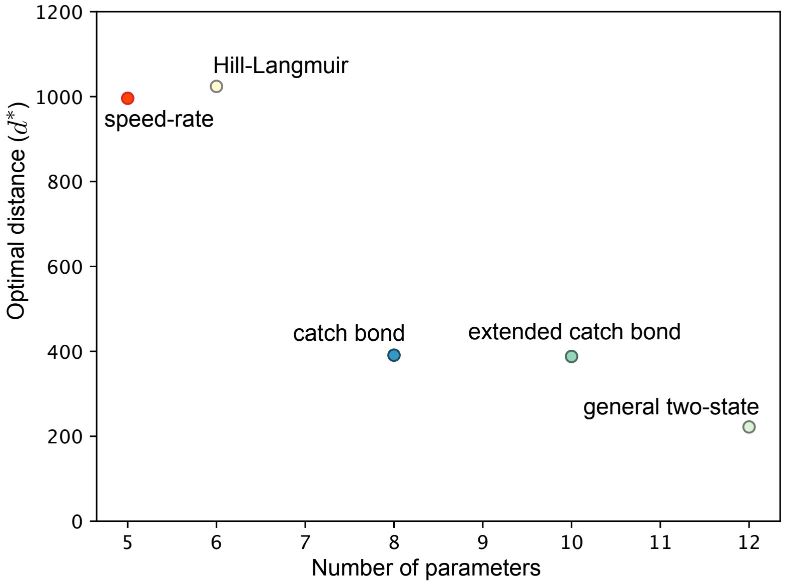


**Figure 4:** Relationship between the number of parameters and optimal distances for the different models studied.

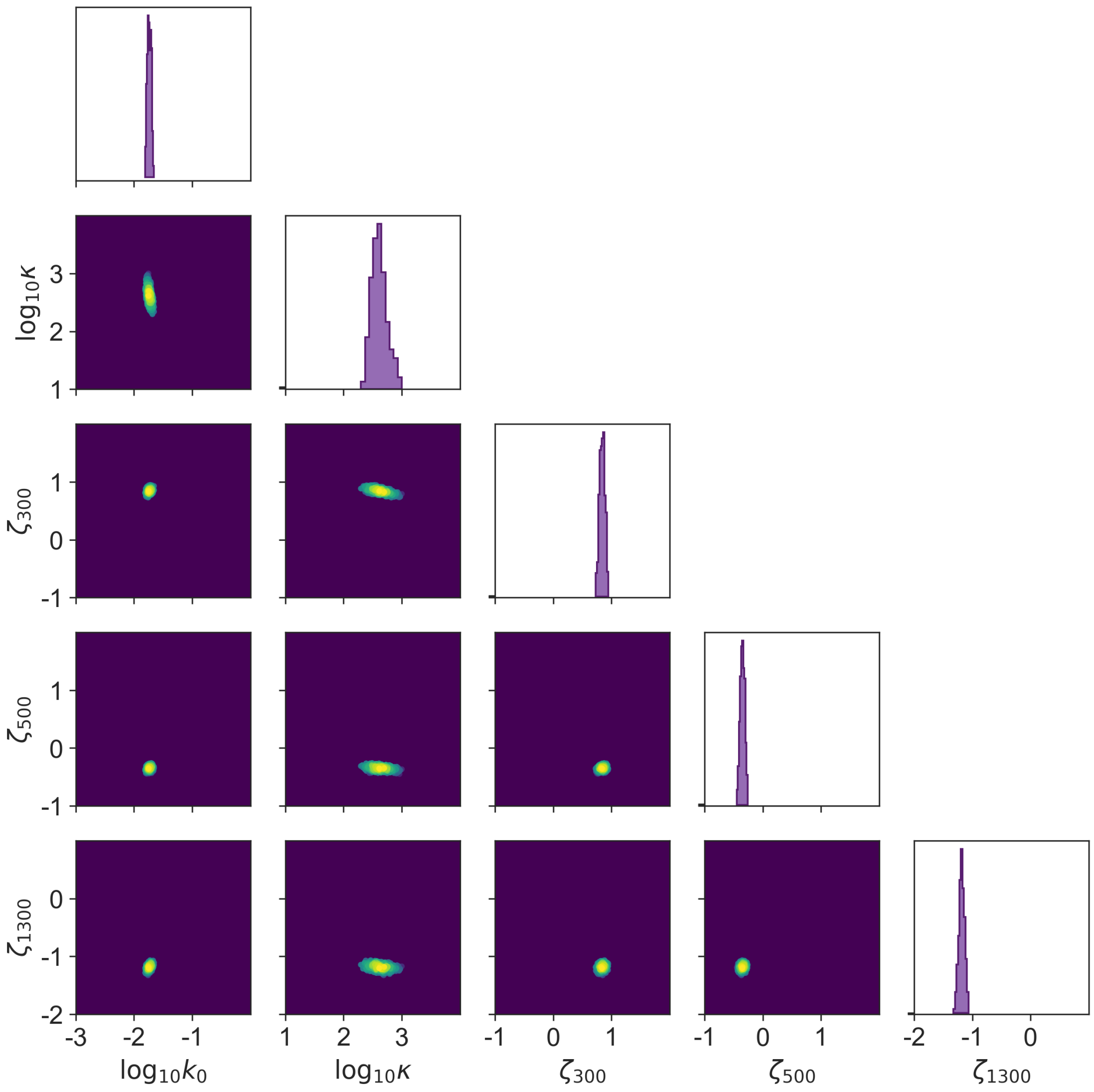


**Figure 5:** Sample of the posterior distribution for the speed-rate model. Marginal posterior (diagonal) and pairwise posterior distributions (off-diagonal), color indicates density of the joint posterior distribution. Units of the rates are in $s^{-1}$. Parameter $\zeta \equiv \beta(\mu_T - \varepsilon)$ indicates the free energy change term in the speed-rate model. Axes ranges correspond with the range of the uniform prior distributions.

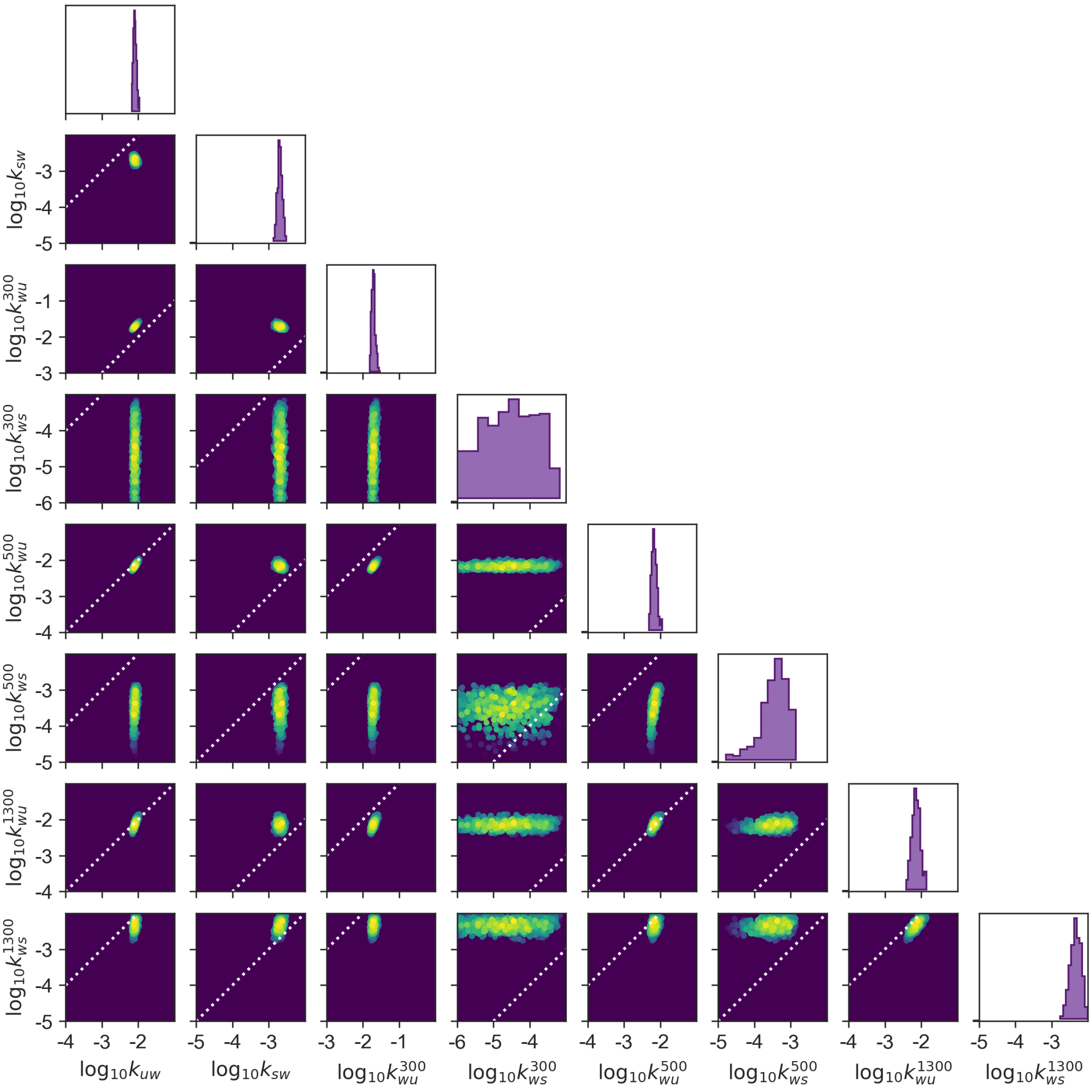


**Figure 6:** Sample of the posterior distribution for the two-state catch-bond model. Marginal posterior (diagonal) and pairwise posterior distributions (off-diagonal), color indicates density of the joint posterior distribution. Units of the rates are in $s^{-1}$. Axes ranges correspond with the range of the uniform prior distributions. Two rates are identical along the guiding dashed lines.

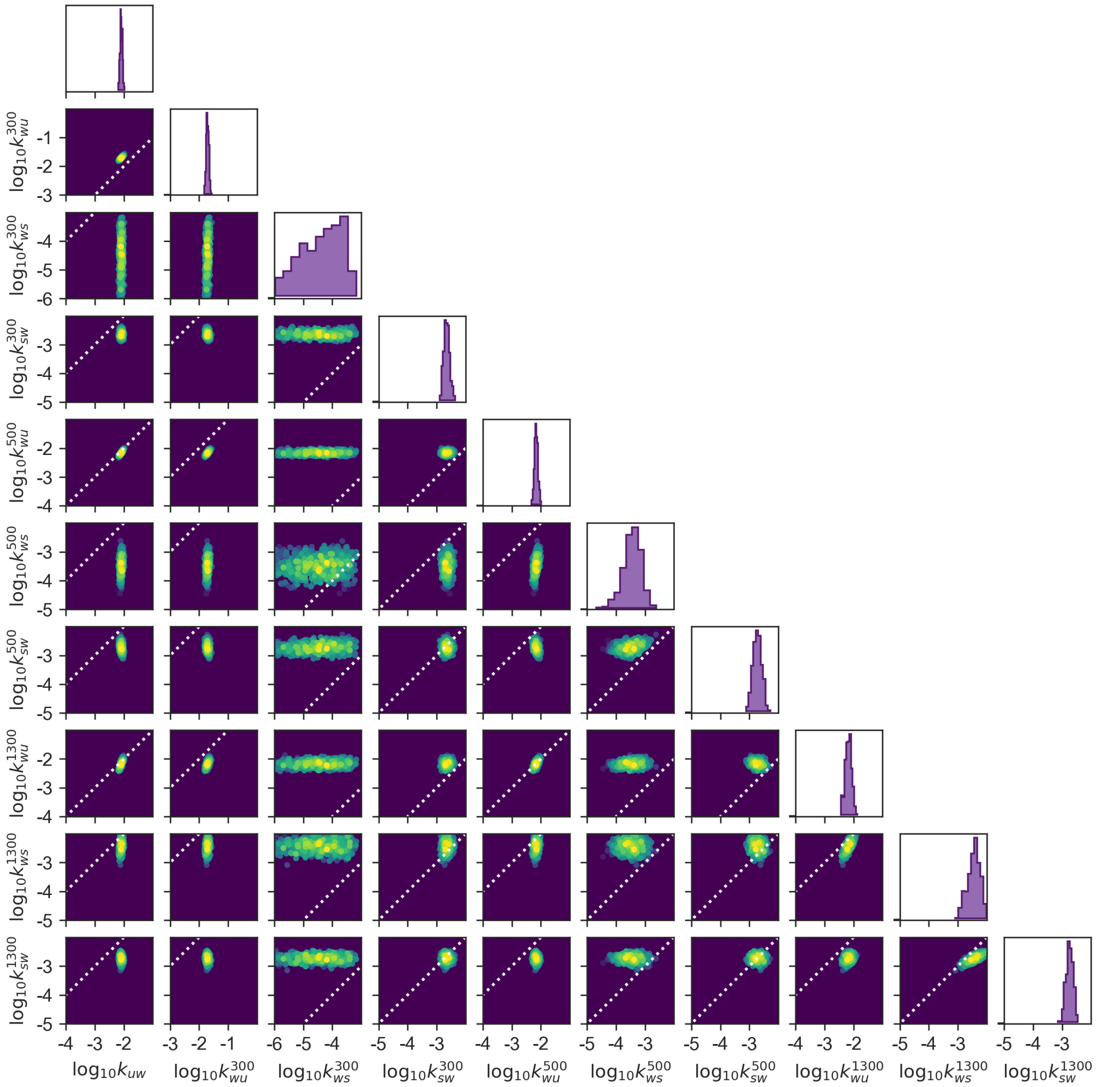


**Figure 7:** Sample of the posterior distribution for the extended two-state catch bond model. Marginal posterior (diagonal) and pairwise posterior distributions (off-diagonal), color indicates density of the joint posterior distribution. Units of the rates are in $s^{-1}$. Axes ranges correspond with the range of the uniform prior distributions. Two rates are identical along the guiding dashed lines.

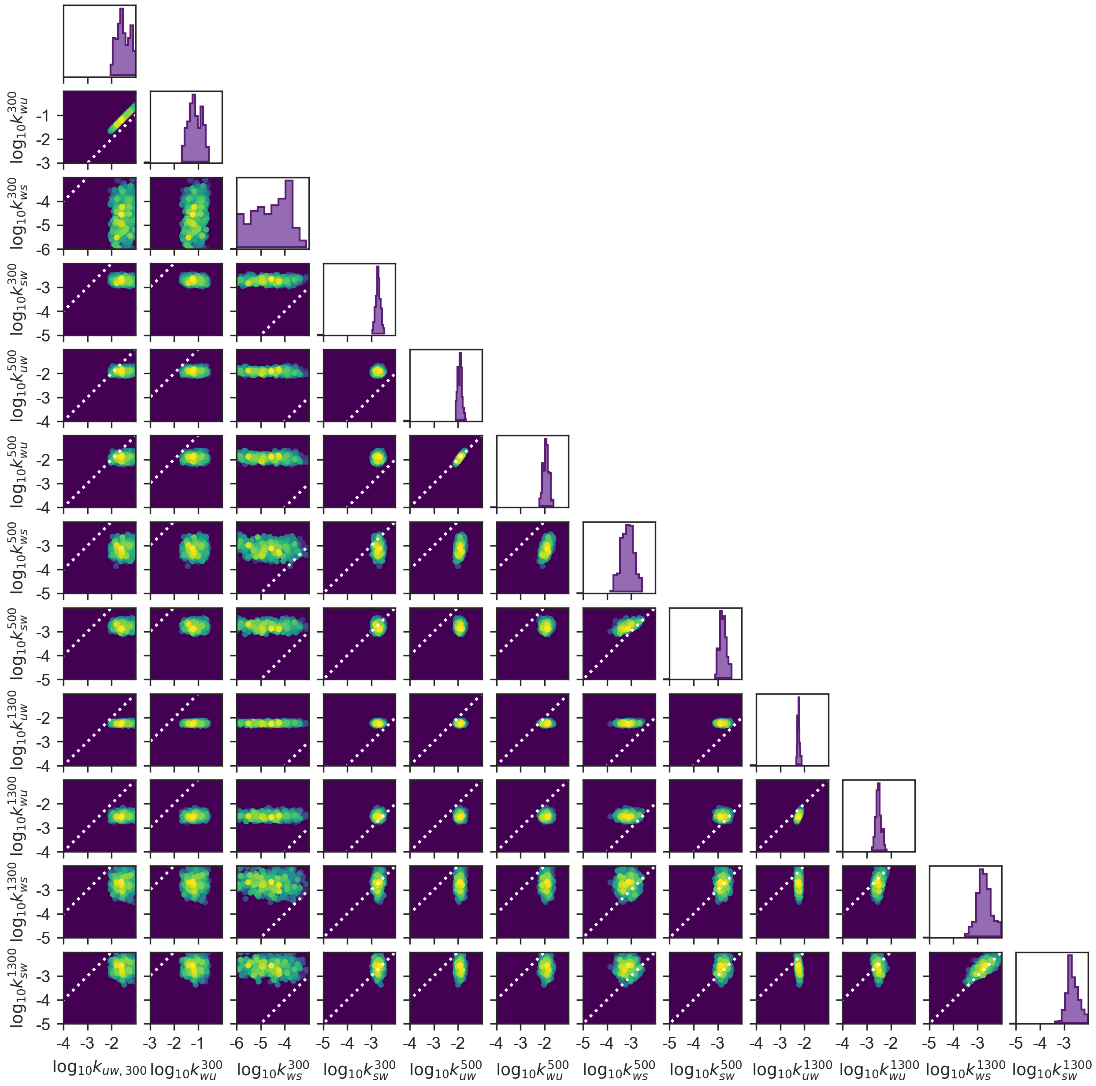


**Figure 8:** Sample of the posterior distribution for the general two-state model. Marginal posterior (diagonal) and pairwise posterior distributions (off-diagonal), color indicates density of the joint posterior distribution. Units of the rates are in $s^{-1}$. Axes ranges correspond with the range of the uniform prior distributions. Two rates are identical along the guiding dashed lines.
